# Modeling Glioma Oncostreams In Vitro: Spatiotemporal Dynamics of their Formation, Stability, and Disassembly

**DOI:** 10.1101/2023.12.14.571722

**Authors:** Syed M. Faisal, Jarred E Clewner, Brooklyn Stack, Maria L. Varela, Andrea Comba, Grace Abbud, Sebastien Motsch, Maria G. Castro, Pedro R. Lowenstein

## Abstract

Glioblastoma (GBM), known for its invasive nature, remains a challenge in clinical oncology due to its poor prognosis. Only 5% of patients live past 2 years. The extensive intra-tumoral heterogeneity, combined with aggressive infiltration into surrounding healthy brain tissue limits complete resection and reduces the efficiency of therapeutic interventions. In previous studies using *ex-vivo* 3D explants and *in-vivo* intravital imaging, we discovered the existence of oncostreams. Oncostreams are accumulations of nematically aligned elongated spindle-like cells constituted by both tumor and non-tumor cells. We observed a direct correlation between the density of oncostreams and glioma aggressiveness, in genetically engineered mouse glioma models, in high-grade human gliomas, and especially in gliosarcomas. Oncostreams play a pivotal role in the intra-tumoral distribution of both tumoral and non-tumoral cells, potentially facilitating collective invasion of neighboring healthy brain tissue. We further identified a unique molecular signature intrinsic to oncostreams, with a prominent overexpression of COL1A1, MMP9, ADAMts2, and ACTA2 - pivotal genes influencing glioma’s mesenchymal transformation and potential determinants of tumor malignancy. COL1A1 inhibition in genetic mouse gliomas resulted in the elimination of oncostreams and induced significant changes in the tumor microenvironment, a reduction in mesenchymal-associated gene expression, and prolonged animal survival. Based on this foundation, we endeavored to model glioma oncostreams *in vitro*, evaluating the potential of various pharmacologic agents on the formation and organization of oncostreams. Using an optimized workflow, oncostreams were established using GFP^+^ NPA cells (NRAs\shP53\shATRX) derived from a genetically engineered mouse model utilizing the Sleeping Beauty transposon system. In-depth global and localized statistical analysis employing Julia programming and R Studio based in-house scripts provided insights into the behavior and organization of glioma cells. Our *in vitro* model led us to probe the impact of factors like cell density, cell morphology, collagen coating, exposure to neurotransmitter agonists, and changes in calcium levels. We also explored interventions targeting specific cytoskeleton structures like non-muscle myosin II B and C, myosin, actomyosin, and microtubules on oncostream formation and organization. In conclusion, our data provide novel information on patterns of glioma migration, which will inform mechanisms of glioma collective invasion in vivo. Through quantitative analysis of these pathologically aggressive and invasive structures, we highlight the importance of potential anti-invasion targets in improving outcomes for GBM patients. Integrating anti-invasive molecules with conventional treatments could significantly enhance clinical benefits.

**Graphical Abstract:** Dynamics of oncostream structure and cellular motility modulation.
This graphical abstract represents the intricacies of the oncostream structure, a proposed model for the collective migration of cancer cells. The central diagram illustrates the oncostream structure, delineated by various treatment conditions radiating outward. Each segment displays a fluorescent micrograph showing the effect of specific inhibitors and compounds on cellular oncostream structure. The array of compounds, including TC-I-15 (α2β1 integrin inhibitor), Collagenase, p-nitro Blebbistatin, Cytochalasin-D, BAPTA-AM, Histamine, Glutamate, 4-Hydroxy acetophenone (4-HAP), Rho-Inhibitor, and Rho-Activator I, are marked on each corresponding segment. Quantitative measures of cellular migration speed, expressed in micrometers per hour (μm/h) are noted for each treatment. Notably, the top half of the diagram reveals the oncostreams’ sensitivity to pharmacological drug treatments, whereas the bottom half shows resistance to these treated conditions. This representation emphasizes the selective effects of pharmacological agents on cancer cell motility within the oncostream framework.

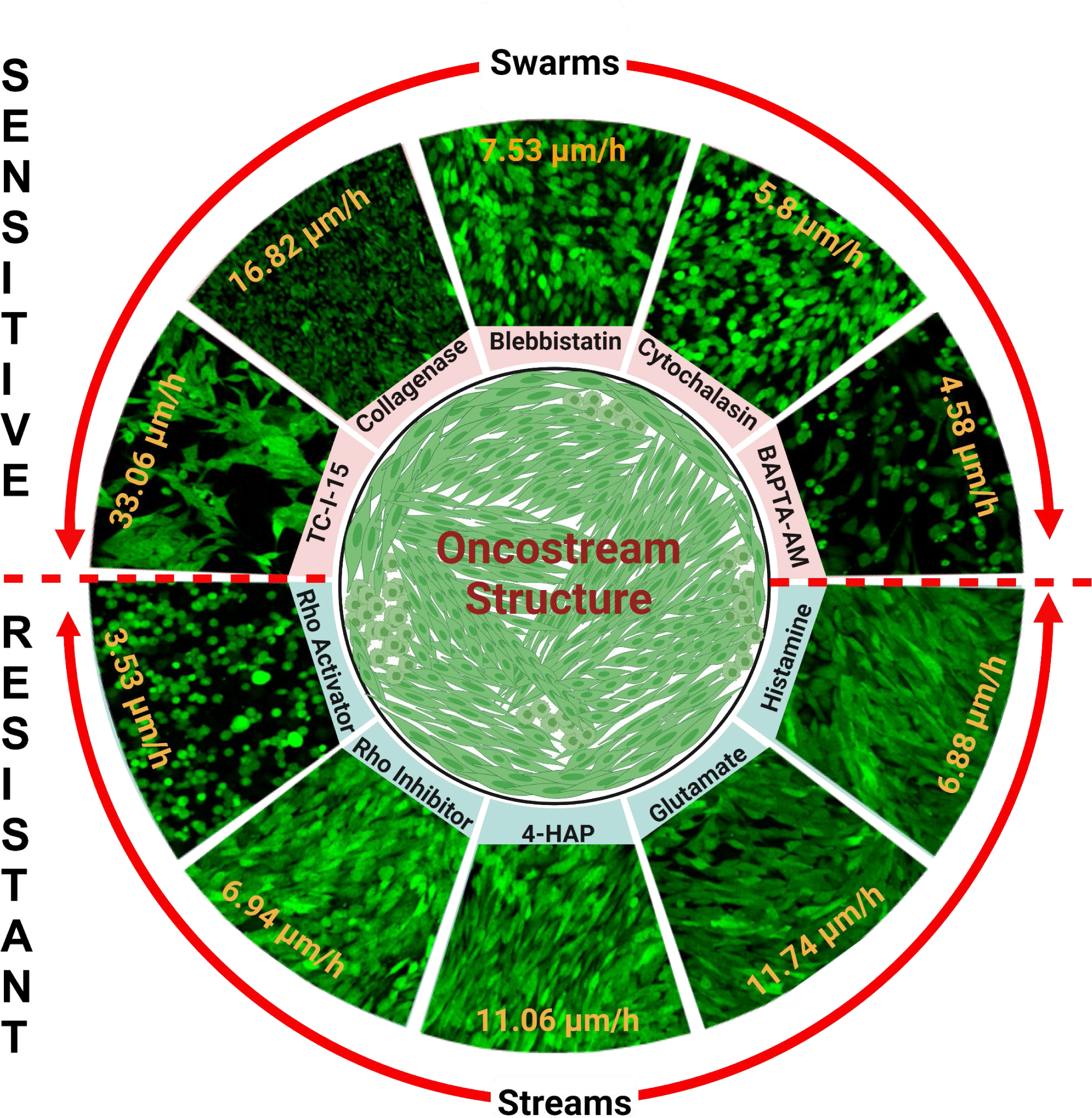

## Introduction

Glioblastoma (GBM) is the most aggressive brain cancer and remains the cancer with the worst prognosis and shortest life expectancy.(*1, 2*) GBM kills through direct invasion of surrounding normal brain tissue rather than metastasizing to distant organs.(*3-5*) Understanding the mechanisms of GBM growth and migration is paramount for future successful treatments. Gliomas extensive infiltration into healthy brain tissue is worsened by significant intra-tumoral heterogeneity, making it particularly resistant to traditional therapeutic modalities and surgical interventions.(*6-9*) Even with the progress of advanced surgical and imaging techniques, surgical resection alone cannot remove all glioma cells, tumors always recur, carrying a very poor prognoses for patients.(*10-13*) The standard of care, involving surgical resection, radiotherapy, and chemotherapy, has made little progress in improving long-term survival over the past two decades.(*14-17*) In the United States, Tumor-Treating Fields (TTFs) have been introduced as an optional treatment for GBM, demonstrated improved progression-free and overall survival in a phase III trial (NCT00916409).(*18, 19*) This highlights the importance for a deeper, more refined understanding of the complex spatiotemporal glioma dynamics.

Recent advancements in gene therapy have shown promise in extending survival rates for glioma, yet tumor relapse remains a significant challenge due to residual invasive cells.(*20-23*) Our recent phase I clinical trial, which combined cytotoxic and immune-stimulatory gene therapy with standard of care (ClinicalTrials.gov identifier: NCT01811992), highlights the importance of addressing this issue.(*22, 24, 25*) In the future phases of our trial, we aim to introduce a specific pharmacological molecule targeting glioma invasion. This approach is anticipated to significantly enhance clinical outcomes by preemptively addressing tumor invasion, potentially transforming the therapeutic landscape for this devastating disease.

We have recently demonstrated the existence and role of oncostreams. Utilizing state-of-the-art ex-vivo 3D explants and *in-vivo* intravital imaging techniques, our group has previously identified a direct correlation between oncostream density and glioma aggressiveness.(*26-28*) We observed this correlation in genetically engineered mouse glioma models and high-grade human gliomas.(*26, 28*) Furthermore, oncostreams have been recognized as pivotal players in the intra-tumoral distribution of both neoplastic and non-neoplastic cells, suggesting a crucial role in the collective invasion of adjacent healthy brain tissues.(*26, 28, 29*) At the molecular level, a specific molecular signature, dominated by the overexpression of COL1A1, MMP9, ADAMts2, and ACTA2 genes, was detected. Deletion experiments demonstrated that COL1A1 is a critical determinant of oncostream formation, and glioma malignancy.(*26*)

Building on this molecular and cellular foundation, we wished to establish an *in vitro* model of glioma oncostreams. We have worked towards replicating these structures *in vitro* to understand their behavior and responses to various pharmacologic agents. Through an optimized workflow, oncostreams were developed *in vitro* using GFP^+^ NPA cells, derived from a genetically engineered mouse model induced using the Sleeping Beauty transposon system. Our comprehensive analytical framework, encoded by Julia and R Studio, provided detailed insights into the multifaceted behavior and organization of glioma cells. We assessed the activities of potential modulators of oncostream structure, including but not limited to cell density, morphological attributes, collagen interactions, neurotransmitter exposure, calcium level variations, and interventions targeting key cytoskeletal components. By providing an integrative understanding of the potential role of oncostreams in glioma invasion, we aim to bridge the gap between spatiotemporal glioma dynamics and potential therapeutic interventions, and thus contribute significantly to the ongoing efforts in improving advancing glioma treatments and the outcomes for GBM patients.

## Materials and Methods

### Animals

All animal studies were conducted in accordance with the Institutional Animal Care and Use Committee (IACUC) of the University of Michigan, referencing protocols PRO00009578, PRO00009599, and PRO00009591. All animals were housed in an AAALAC-accredited animal facility and were observed daily. Our research included both sexes, and we used the C57BL/6 mouse strain procured from The Jackson Laboratory (strain no. 000664).

### Hematoxylin and Eosin (H&E) staining

This process begins with deparaffinization and hydration, using xylene and graded concentrations of ethyl alcohol. Initially, the sections are treated thrice with xylene for 5 minutes each, followed by two 2-minute treatments each with 100%, 95%, and 70% ethyl alcohol, and finally rinsed in water for 2 minutes. The staining phase involves immersing the slides in hematoxylin for 1 minute and 30 seconds, then rinsing them under running tap water until clear. This is followed by a quick dip in the bluing solution, and another 2-minute rinse under running tap water. The slides are then treated with 80% ethyl alcohol for 2 minutes and stained with eosin for 5 minutes. In the dehydration stage, the slides are first dipped 3-4 times in 80% ethyl alcohol to rinse out excess eosin, followed by two 2-minute treatments each in 95% and 100% ethyl alcohol. Finally, the slides undergo three 3-minute treatments in xylene. Cover slips were applied using xylene-based mounting medium. This staining ensures that chromatin in nuclei is stained dark blue (by Hematoxylin), while the cytoplasm is stained pink (by Eosin).

### H&E and Immunofluorescence (IF) Imaging

Mouse brain samples were perfused with Tyrode solution followed by PFA, then post-fixed in 4% PFA for 48 hours at 4°C, and paraffin-embedded at the University of Michigan’s facility using Leica ASP 300 and Tissue-Tek.(*26, 28*) Sections (5 μm) were obtained using a Leica rotary microtome. For histopathological examination, sections underwent Hematoxylin and Eosin (H&E) staining. Concurrently, DAPI staining was performed for Immunofluorescence (IF) in GFP-positive cells. Imaging of H&E-stained sections utilized an Olympus BX53 Upright Microscope, selecting ten random fields per section to encompass tumor heterogeneity. For immunofluorescence, paraffin-fixed glass coverslips post oncostreams formation was imaged using a Zeiss LSM 880 laser scanning confocal microscope. The resulting data were analyzed using the Zeiss Zen (Blue edition) software, version 2.5.

### Generation of GFP^+^ NPA glioma cells derived from the genetically engineered mouse glioma model (GEMM)

In our lab, we have developed a unique approach to model glioma by introducing oncogenic plasmid DNA into the neural stem cells of postnatal mouse brains, utilizing the Sleeping Beauty (SB) transposon system. We employed fully verified plasmid sequences for this approach. The specific plasmid sequences used to generate the NPA GFP^+^ tumors were: (1) pT2C-LucPGK-SB100X for dual transposon and luciferase expression, (2) pT2-NRAS-G12V to instigate RTK/RAS/PI3K pathways that drive oncogenesis, (3) pT2-shp53-GFP4 for attenuating p53 to knock-down tumor suppressor p53 expression, and (4) pT2-shATRx-GFP4 for ATRX knock-down. Notably, the pT2CAG-NRAS-G12V and pT2-shp53-GFP4 plasmids were a kind contribution from the Ohlfest laboratory at the University of Minnesota. The methodologies employed for glioma induction, as well as the generation of glioma cells, have been comprehensively detailed by our team in prior publications including Calinescu et al., Koschman et al., Nunez et al., Alghamri et al., and Comba et al.(*30-34*)

#### Neonatal plasmid delivery

Plasmids were injected peri-ventricularly into both female and male C57BL/6 mice at postnatal day 1 (P01). The plasmid mixture included: SB/Luc, NRAS^G12V^, shp53, and shATRX, in mass ratios of (1:2:2:2). 20μg of the plasmids were mixed into a 40μL solution, combined with *in vivo*-jet PEI® (Polyplus Transfection, New York NY, 201-50G) (3.4μL per 40μL plasmid mixture) and 10% glucose, resulting in a final solution with a 0.5 µg/μL DNA concentration. The plasmid solution was kept at room temperature for a minimum of 20 minutes prior to injection. Neonatal mice were anesthetized (via hypothermia) by placing them on wet ice for 2 min, while maintaining a body temperature between 2-8°C per ULAM guidelines. Subsequently, neonatal mice were positioned on a temperature-controlled neonatal stereotaxic frame cooled to 2-8°C to maintain anesthesia for the remainder of the procedure. The injection point was determined using the coordinates set to 1.5 mm AP, 0.8 mm lateral to the lambda, and 1.5 mm deep to inject 0.75μL plasmid mixture at a rate of 0.5μL/min into lateral to the lateral ventricle of neonatal mice as described previously by us.(*32*)

#### Monitoring tumor development and growth through bioluminescence

To monitor the plasmids’ presence and uptake by the neonatal mice, we utilized *in vivo* bioluminescence measured using the IVIS® Spectrum imaging system (Perkin Elmer, Waltham MA). One day after plasmid injection, 30μL of Luciferin (30 mg/mL) was injected subcutaneously into each pup. We then imaged the pups on an IVIS® Spectrum imaging system, employing settings like automatic exposure, large binning, and aperture f =1. Any pups that did not exhibit luminescence were humanely euthanized according to approved procedures. In the adult mice, we carried out daily health checks for signs of morbidity as described before.(*30-32, 34, 35*) The growth of the tumor was periodically observed every 15 days utilizing the IVIS® Spectrum imaging system. For this, a 100 µl dose of 30 mg/ml Luciferin solution was administered intra-peritoneally using a 1 ml syringe fitted with a 26-gauge needle, consistent with earlier described methods. This methodology allowed us to evaluate the progression of the tumor. When adult mice began showing signs of morbidity, we proceeded with transcardial perfusion-fixation, as previously described. Brain tumors obtained were used for paraffin embedding and posterior sectioning. Additionally, viable non-perfused tumor tissue was utilized for the generation of primary neurospheres.

### Generation of primary neurospheres from SB tumors

From mice bearing tumors via the SB approach, we selected a mouse with a substantial tumor. This was verified using an in vivo imaging system, with bioluminescence readings between 10^7^-10^8^ photons/s/cm2/sr. Mice were euthanized with isoflurane anesthetic, decapitated, and their brains were carefully harvested on ice. The tumor mass (visible by the green fluorescence) was dissected with forceps and scalpel under a fluorescence dissection microscope. The isolated tumor tissue was then placed in 1mL of neural stem cell (NSC) media and subjected to mechanical homogenization using a plastic pestle as described previously.(*26, 31, 32, 35, 36*) This NSC media comprised DMEM/F12 with L-Glutamine (Gibco®, 11320-033), supplemented with 1% B27 (Gibco®, 12587-010), 1% N2 (Gibco®, 17502-048), Penicillin-Streptomycin (100 U/mL) (Corning, Cellgro®, Corning NY, 30-001-CI), and Normocin™ (1X) (InvivoGen, ANT-NR-1). Tissue was then digested with 1 mL of enzyme-free dissociation agent HyClone™ (GE Healthcare Life Sciences) for 5 min at 37°C. The tissue suspension was then filtered through a 70 µm cell strainer, rinsed with an additional 10 mL of media, centrifuged at 300 g for 4 min, and the supernatant was removed. The remaining cell pellet was resuspended in fresh NSC media and seeded into a T25 culture flask. After incubating for three days at 37°C (95% air and 5% CO2), the formation of robust, free-floating neurospheres was observed. Cell cultures were maintained at 37°C with 95% air and 5% CO2. Additionally, hFGF and hEGF (ThermoFisher Scientific, AF-100-18B-1 and AF-100-15-1, respectively) were supplemented twice per week at 20 ng/ml.

### Adapting floating neurospheres to an adherent phenotype

To transition these primary NPA GFP^+^ neurospheres to an adherent phenotype, we first coated T25 flasks with 15µg laminin in 3 ml of DPBS, enriched with CaCl_2_ and MgCl_2_ (Sigma, D8662-500ML). Flasks were then incubated for two hours at 37 °C under 5% CO_2_ conditions. Following this, we transferred the coated flasks to 4 °C to solidify overnight. Prior to introducing the glioma cells, it’s essential to rinse the flasks twice with cold DPBS enriched with CaCl_2_ and MgCl_2_ to remove unbound laminin followed by one washing with DPBS without CaCl_2_ and MgCl_2_. NPA GFP^+^ neurospheres were then treated with Accutase® cell detachment solution (Biolegend, 423201) for 2 min to make a single cell suspension. This single cell suspension was then cultured on the laminin coated T25 flasks using DMEM medium containing 10% FBS. Notably, we ensured the cells underwent at least 5 passages before setting up any time-lapse experiments. After these passages, we observed that the NPA GFP^+^ cells had begun to show collective organization patterns, particularly structures resembling oncostreams.

### Preparation of glass bottom culture dishes for the development of oncostreams *in vitro*

We have optimized a workflow for preparing glass-bottom culture dishes conducive to the development of oncostreams *in vitro*, as depicted in **Figure 2C**. Initially, we acid-washed the 35mm dishes (Lot#202001; Ibidi; 81218-200) with 1M HCl at 60°C to bolster polypeptide adherence, followed by rinses with milli-Q water and an ethanol treatment. The dishes were then coated with a Poly-D-lysine solution (100μg/ml) diluted in DPBS (pH 7.4) overnight at 4°C. On the next day, after drying and rinsing, laminin coating was applied, prepared from a thawed stock, and diluted in DPBS enriched with CaCl_2_ and MgCl_2_ to 10μg/ml. Following laminin-priming, the dishes were incubated at 37°C for 1-2 hours to facilitate binding and then stored at 4°C to solidify overnight. Prior to cell seeding, dishes were rinsed with cold PBS enriched with CaCl_2_ and MgCl_2_. NPA GFP+ cells, adapted to an adherent phenotype, were then seeded onto the laminin-coated dishes in DMEM medium with 10% FBS. After 2 hours, 1 ml of fresh medium was added. The final step involved an 18h incubation to ensure thorough cell adherence to the glass surface.

### Understanding tumor cell dynamics through post-imaging analysis

To gain a deeper understanding of the complex invasion and infiltration processes during GBM progression, we developed an *in vitro* model that simulates the *in vivo* behavior of glioma invasion. This model aids in identifying pharmacological agents, which eventually target unresectable invasive cells before surgery. We (SM) developed a detailed statistical methodology to analyze glioma growth patterns,(*26, 28*) allowing the identification of pharmacologic agents that disrupt oncostreams effectively *in vitro*. Our statistical analysis involves the determination of cell trajectories, path filtering, speed distributions, angular velocities, velocity vectors, likelihood analysis, pairwise and nematic correlations, and relative position.(*26, 28*)

### Data Collection via time-lapse imaging

We utilized a laser scanning confocal microscope, the LSM 880 with Axio Observer from Zeiss, Germany, which was synchronized with the Zen (Blue edition) version 2.5 software. This setup was used to acquire and process time-lapse videos. The time-lapse videos were taken at intervals of 10, 5 and 1.5 minutes, and were captured using a 20X lens encompassing areas of 1190.60 x 1190.16 µm, 850.19 x 850.19 µm, and 425.10 x 425.10 µm, with each pixel measuring 0.415 x 0.415 µm. For further statistical analysis, we utilized tools such as the TrackMate plugin, Image J Software, Julia, and RStudio. We ran the respective scripts that are available by us on GitHub and Zenodo (https://zenodo.org/records/7587235). Detailed instructions for evaluating glioma cell dynamics are provided below:

### Cell tracking via Fiji plugin “TrackMate”

For tracking cell migration, we utilized Fiji software (<u>Fiji (imagej.net)</u>) integrated with the TrackMate plugin (<u>TrackMate (imagej.net)</u>). Once the software was activated, the initial panel displayed detailed image dimensions. Proceeding with “Next” introduced a secondary panel where the size of the cell, referred to as “blob” could be specified. We selected a size of 20 µm, set a threshold of 1, and employed the Difference of Gaussian (DoG) detection method. As we progressed and the loading bar filled up, the interface transitioned to the tracking settings. At this juncture, we chose the “Simple LAP” tracker. Following this, the “Track filter” interface allowed us to refine the tracking data. Ideally, for every marked area in the image, we retained tracks that had a minimum of ten pathways and removed any that didn’t meet this criterion. As we delved deeper into the analysis of each time-lapse video, various cell pathways were observed. By navigating to the “Analysis” section, three detailed windows were presented: “Spots”, “Tracks Statistics”, and “Link in Track Statistics”. For comprehensive analysis, the CSV files labeled “Spots” and “Tracks Statistics” are the essential ones.

### Visualization of trajectories using Julia programming

This step dealt with extracting meaningful trajectories patterns from the raw data. Using the CSV outputs named *spots.csv and *tracks.csv from the previous step, we visualized cell positions and their movement trajectories. For those who are interested in practical examples of these files, a repository can be found on Zenodo (https://zenodo.org/records/7587235) under the directory ‘data tracking step1_data_trackMate/NPA_stich_3’. By executing the designated Julia script, comprehensive cell trajectories over the course of the time-lapse datasets were generated, as depicted in several Figures as *cell tracking*.

### Trajectories filtering process

To address the erratic behavior often evident in cell trajectories, we used a filtering Julia script. Using the data previously derived from TrackMate, we smoothed the trajectories, allowing us to estimate the velocity of each cell at specific time points. We employed a Gaussian kernel filter with a standard deviation of σ = 2 and a stencil of 9 points to achieve this. This step was executed via a Julia script, specifically named step2_filter_data.jl, which can be found in the ‘SRC’ folder on our Zenodo repository (https://zenodo.org/records/7587235). Once the script was ran, the trajectory graphs of the time-lapse datasets were refined, as illustrated in several Figures named as “*filtered path”*. This filtration step also enabled us to determine the positions Xi(t) and velocities Vi(t) of each cell *(i)*. Additionally, from the velocity Vi(t), we deduced the velocity direction θi(t), setting the stage for a more in-depth understanding of the migration pattern of the cells. The resultant data showcased a path derived from previous step (cell tracking) and a filtered path. This smoother path facilitated the estimation of cell velocity at any given time step. All refined trajectories were stored in a jld2 file within the ‘data_tracking’ folder on our Zenobo repository, labeled as step2_data_filtered.

### Velocity and correlation distribution analysis

At this juncture, each cell was marked by its distinct position *x_i_* ∈ R^2^ and an associated velocity *v_i_* ∈ R^2^. To extract more profound insights, we separated the velocity vector **(v)** into its magnitude – the speed (scalar) c = |**v**| and its directional angle θ, depicted as v = c (cos *θ*, sin *θ*). Firstly, we assessed the velocity angle distribution, which provided the predominant direction in which the cells moved. Based on the visible structures (Oncostremas) within the *in vitro* culture at the movie’s conclusion, we manually designated distinct zones for closer inspection; for a clearer picture, please refer to **Supplementary Figure S3A-B**. To pinpoint each zone’s exact coordinates, we employed the Fiji coordinate tool, which revealed the cursor’s x and y placements in µm. These pinpointed coordinates, crucial for our zone-specific evaluations, were diligently archived in a JSON format, properly named “coordinates.json”. A sample of this file is accessible on our Zenodo repository (https://zenodo.org/records/7587235). Using file named “coordinates.json”, Julia software demarcated the individual zones, employing the script “step3_zone_create_df.jl”, which is available on Zenodo repository. Launching this script resulted in the creation of data frames, each detailing the analytical breakdown of the designated zones. For a comprehensive analysis, it was essential to initiate the script twice, focusing on velocity for one run and correlation for the other. A minor tweak in Line 13 of the script allowed this switch between “velocity” and “correlation”. Subsequent visual portrayals for either velocity or correlation were crafted via specific Julia scripts, listed on Zenodo. Our exhaustive findings, including datasets on angle velocity, speed metrics, and velocity heatmaps per zone, reside within the df_velocity directory. Concurrently, insights on correlations and relative positions are cataloged in the df_correlation directory. A comprehensive visual representation of these datasets is hosted on Zenodo repository (https://zenodo.org/records/7587235).

### Classification of collective migration patterns using RStudio-driven likelihood analysis

To discern specific migration patterns within each designated or global zone--be it flock, stream, or swarm--we meticulously analyzed the angle *θ_i_* within a designated zone. Three distinct distributions were employed for this purpose: the (wrapped) Gaussian which hinted at a *flock* pattern, the symmetrized Gaussian which leaned towards a *stream*, and the constant function symbolizing a *swarm*. Our methodological approach hinged on the *Akaike weight* (AW) distribution analysis, which effectively determined the predominant distribution for *θ_i._* For a streamlined evaluation and to pin down the precise character of the zone or global movie, we employed a dedicated R script, accessible via Zenodo repository. This script, once executed, resulted in a visual representation—a zone distribution graph—in R Studio, accompanied by the consequential Akaike weight outcomes. It’s essential to highlight that an AW approximating 1 was considered as the most indicative, emphasizing the highest likelihood, and any significant deviations from this value were typically disregarded.

### Influence of cell density on glioma self-organization

To elucidate the impact of cell density on glioma self-organization, we systematically varied cell seeding densities. These densities ranged from a low density of 1 x 10^5^ cells to a high density of 2 x 10^5^ cells. After 18h of incubation at 37 °C under 5% CO_2_ conditions, the time-lapse video were acquired at 5-minute intervals, using a 20X lens encompassing areas of 1190.60 x 1190.16 µm with each pixel measuring 0.415 x 0.415 µm in width and height. The time-lapse raw data was processed as described in section “data collection via time-lapse imaging”. Our aim was to determine any potential correlation between cell density and the formation of oncostreams. The specific experimental procedures employed, as well as the methods of data analysis are elaborated in the “Statistical Analysis” section.

### Collagenase treatment

To assess the influence of extracellular matrix components on glioma behavior, we examined the effect of collagenase treatment on oncostream formation. Glioma cells were seeded at a high density (2 x 10^5^), targeting the observation of the treatment’s impact on oncostream dismantling. Cells were allowed to self-organize and initiate oncostreams over a 36-hour period post-seeding. After this, they were treated with a 15 U/ml of collagenase enzyme, aiming to degrade the surrounding matrix without inducing significant cell death. Time-lapse videos were subsequently set up to capture images at 10-minute intervals, utilizing a 20X objective which covered fields of 1190.60 x 1190.16 µm, with individual pixels measuring 0.415 x 0.415 µm in both width and height. The raw data acquired from the time-lapse imaging was then processed as detailed in the “Data collection via time-lapse imaging” section. Our main objective was to delineate the relationship between matrix degradation induced by collagenase and the propensity of glioma cells to dismantle oncostreams. The explicit data analysis methods are further elaborated in the “Statistical Analysis” section.

### Impact of neurotransmitter agonists and calcium chelation on oncostream integrity

To evaluate the role of neurotransmitters and calcium on the dynamics of oncostream formation, we employed histamine at a concentration of 5μM, glutamate at 1mM, and the calcium chelator BAPTA-AM at 5μM. BAPTA-AM is known to chelate intracellular calcium, thereby potentially modulating cellular behaviors associated with calcium signaling. Glutamate, an excitatory neurotransmitter, plays a multifaceted role in neuronal communication. Unlike most neurotransmitters, glutamate possesses the unique ability to bind to four distinct receptors, underscoring its robust influence in stimulating and facilitating inter-neuronal communication. Following treatment, time-lapse imaging was initiated, capturing images at 10-minute intervals. The field of view spanned an area of 425.10 x 425.10 µm, with each pixel distinctly measuring 0.415 x 0.415 µm in both width and height. This methodology aimed to elucidate the correlation between neurotransmitter activity, calcium chelation, and their collective impact on oncostream disruption dynamics.

### Cytoskeleton-targeting antagonists

To decipher the role of crucial molecular components associated with cell contractility and cytoskeletal organization in oncostream formation, we employed a panel of specific antagonists. These included:

a. *4-hydroxy aceto phenone (4-HAP)*: A compound that targets non-muscle myosin II B and C.(*37*) Used at a concentration of 4 µM, the effects of 4-HAP are slow and develop progressively over time. This necessitates a longer time-lapse interval of 5 minutes, compared to the 90 seconds required under other conditions.
b. *Para nitro blebbistatin (pnBB):* A non-phototoxic, photostable myosin inhibitor used at a concentration of 5 µM, geared towards actomyosin interference. Due to its rapid impact on cells, imaging was set to capture every 90 seconds.
c. *Cytochalasin D:* A cell-permeable and potent inhibitor of actin polymerization. Deployed at a concentration of 5 µM, it was used specifically to disrupt actin filaments. The fast-acting nature of this compound required a capture interval of every 90 seconds.
d. *Nocodazole:* An agent introduced to perturb microtubule dynamics, used at a concentration of 2 µM. It induced swift cellular responses requiring a 90-second imaging interval.

For spatiotemporal analysis, glioma cells were seeded at a density of 2 x 10^5^ on laminin-coated glass-bottom culture dishes. This setup was aimed at closely observing the effect of the treatments on oncostream dismantling. After seeding, the cells were given a 36-hour period to self-organize and initiate oncostream formation. Throughout the investigation, the imaging area was maintained at 425.10 x 425.10 µm, with a pixel resolution of 0.415 x 0.415 µm for both width and height. Our goal was to accurately document the dynamic alterations induced by these pharmacological agents, shedding light on their potential therapeutic applications.

### Inhibition of α_2_β_1_ integrin using TC-I-15

To explore the implications of inhibiting α2β1 integrin in oncostream formation and dynamics, we employed the thiazolidine-modified compound 15 (TC-I-15). TC-I-15 is a selective high-affinity small-molecule inhibitor of integrin α2β1. This compound has been previously shown to allosterically inhibit integrin activation and adhesion of NPA glioma cells to laminin-coated surfaces. For glioma dynamics analysis, glioma cells were seeded on laminin-coated glass-bottom culture dishes. 6 hours post-seeding, TC-I-15 was introduced at a concentration of 5 µM. This setup was strategically chosen to scrutinize its effect on blocking oncostream formation in our in vitro model. Given the compound’s nature and its predicted influence on the cells, we established a time-lapse video capture interval of every 5 minutes. Throughout the confocal microscopy, the imaging area remained consistent at 425.10 x 425.10 µm, ensuring a pixel resolution of 0.415 x 0.415 µm for both dimensions. The overarching objective was to provide a detailed assessment of the dynamic cellular changes elicited by TC-I-15, further advancing our understanding of its therapeutic potential in modulating oncostream dynamics.

### Rho Family GTPases Modulation and its Influence on Cell Behavior

To discern the impact of Rho family GTPases modulation on cellular self-organization and collective invasion, we utilized specific agents for both inhibition and activation. The Rho inhibitor (Cytoskeleton, Inc, Cat# CT04) was administered at a concentration of 2μg/ml, while the Rho activator I (Cytoskeleton, Inc, Cat# CN01) was similarly introduced at 2μg/ml. To monitor the real-time cellular responses to these modulations, we performed confocal microscopy to acquired time-lapse videos at intervals of every five minutes. Throughout these observations, the imaging parameters were maintained with an area of 425.10 x 425.10 µm, and each pixel was distinctly resolved at 0.415 x 0.415 µm in both width and height.

## Results

### Histopathological identification of oncostreams in glioma

Hematoxylin and eosin-stained sections of human primary glioma and a genetically engineered mouse model revealed distinct oncostream structures (**Figure 1**). In both human and mouse gliomas, oncostreams were characterized by densely packed, spindle-like cells, forming elongated structures within the tumor mass as demonstrated before.(*26-29*) Detailed examination showed these oncostreams, outlined with dotted black lines for clarity, throughout the tumors, as well as close to regions of tumor invasion. The presence of round cells was noted outside oncostreams, illustrating the morphologically complex and heterogenous cellular tumor microenvironment.

**Figure 1:**
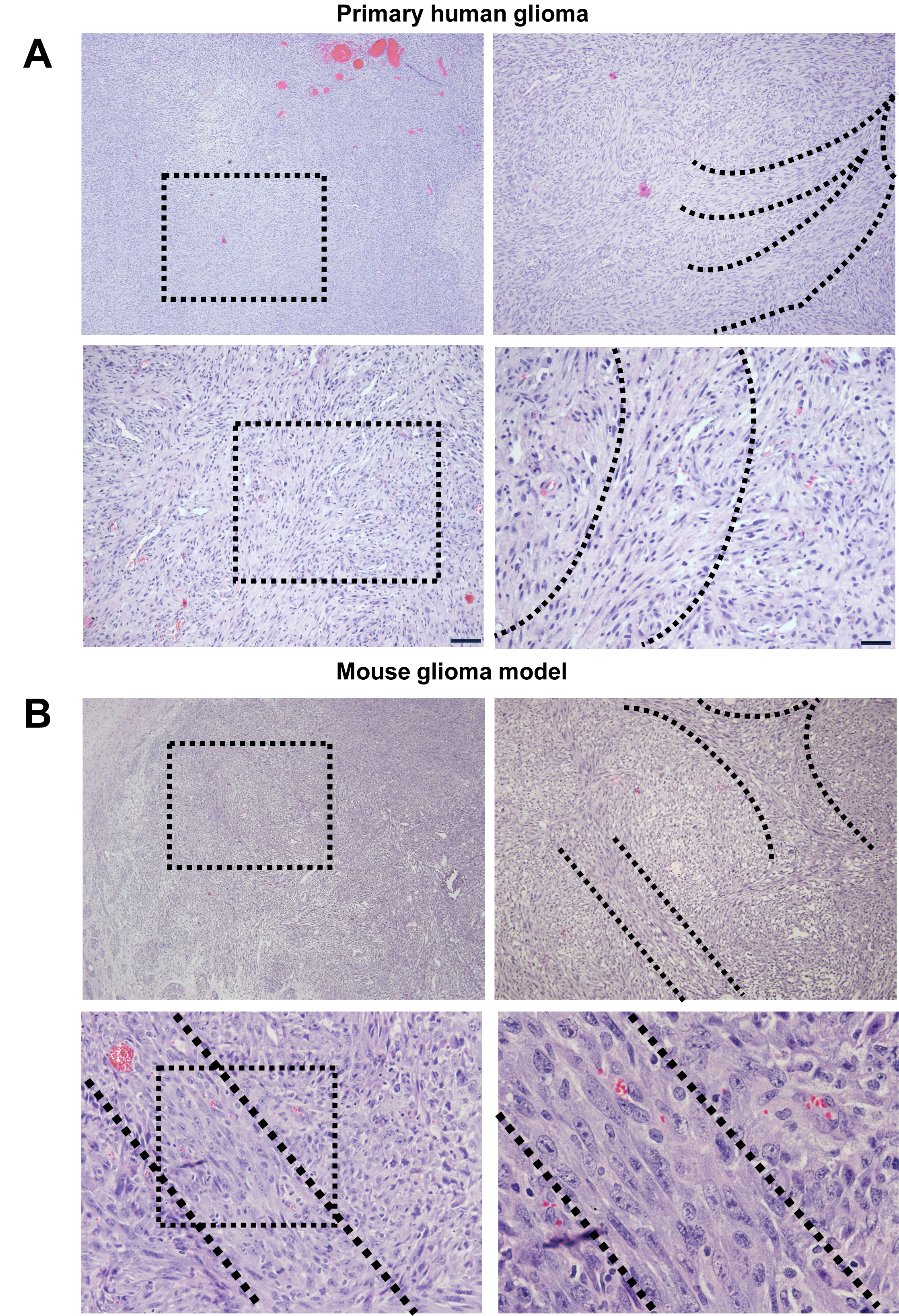
Neuropathological illustration of oncostreams in human and mouse glioma. Hematoxylin and eosin-stained sections displaying oncostreams in human primary glioma **(A)** and a genetically engineered mouse model of glioma **(B).** In both panels, the left panel provides a lower magnification view of the tumor, highlighting a specific region with a black box, which is then shown at higher magnification in the right panel. Oncostreams, identified as bundles of spindle-like cells, are outlined with dotted black lines, demonstrating their path through the tumor. The presence of these structures has been associated with invasive behavior in brain tumors. Round cells are also visible outside the oncostreams.

**Figure 2:**
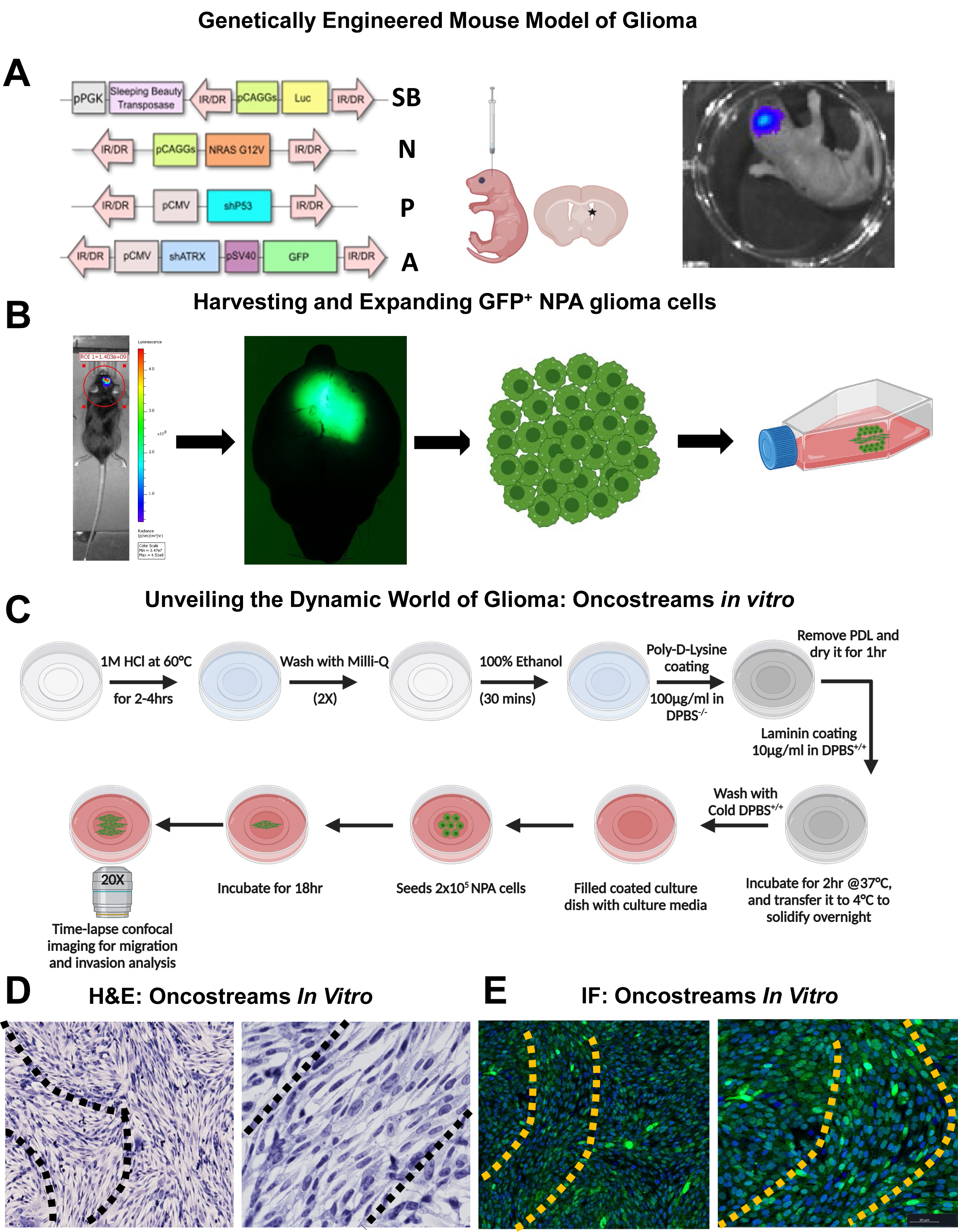
Development and acquisition of oncostreams from GFP^+^ NPA glioma cells *in vitro*. This figure outlines the generation of GFP^+^ NPA high-grade glioma (HGG) in mice and the subsequent development of oncostreams *in vitro*. **(A)** Transposable fragments of the plasmids including NRAS **(N)**, shP53 **(P)**, and shATRX **(A)** used to induce GFP^+^ NPA HGG via the Sleeping Beauty **(SB)** transposon system are shown, which include a luciferase reporter for bioluminescence tracking. **(B)** Detailed procedure for inducing GFP^+^ NPA in neonatal mice: Brain stem cells of neonatal mice are transfected with SB transposase for *in vivo* integration of NPA-inducing genetic alterations into the subventricular zone cells, incorporating GFP expression from the ATRX plasmid. Bioluminescence confirmation via IVIS-Imaging ensures successful transgenesis. **(B)** The emergence of HGG is monitored *in vivo* through luciferase-driven bioluminescence. Upon clinical signs of tumor burden, mice are either perfused for IHC analysis or GFP^+^ NPA cells are isolated from the tumor tissue, identified by ATRX knockdown (KD) green fluorescence, for *in vitro* culture and expansion. **(C)** The step-by-step workflow for in vitro oncostreams development and setting up time-lapse imaging to capture the migration patterns of glioma cells. **(D)** Hematoxylin and eosin-stained images depict oncostreams in cultured NPA glioma cells. **(E)** Immunofluorescence characterization of *in vitro* oncostreams shows elongated nuclei (stained with DAPI in blue) and NPA glioma cells (in green).

### Establishment and characterization of oncostreams in vitro

The GFP^+^ NPA high-grade glioma was successfully induced in mice using a combination of NRAS, shP53, and shATRX plasmids via the Sleeping Beauty transposon system, as illustrated in **Figure 2A**. Oncostreams were developed from isolated GFP^+^ NPA cells cultured *in vitro*, following a methodical process that involved initial tumor induction in neonatal mice, with subsequent isolation and expansion of tumor cells (**Figure 2B**). The *in vitro* model closely replicated the oncostream structures observed *in vivo*.(*26, 27*) Dynamic behaviors of glioma cells during oncostream formation were acquired using time-lapse imaging (**Figure 2C**). ECM components are critical to the organization and migratory behavior of glioma cells in oncostreams. To optimize the workflow for developing oncostreams in vitro, we investigated the impact of laminin on their formation, GFP^+^ NPA glioma cells were cultured without laminin on poly-D-lysine coated dishes. Cells failed to form structured oncostreams over 45 hours, instead cells clustered densely, suggesting a disordered migratory pattern (see **Supplementary Figure S1**). This highlights the pivotal role of laminin as a key ECM component facilitating the structured assembly of oncostreams. Histological and immunofluorescence analyses of cultured NPA glioma cells confirmed the formation of oncostreams **Figure 2D and 2E**. These structures displayed elongated nuclei (H&E, and DAPI-stained) and were primarily composed of NPA glioma cells (GFP-positive), as shown in **Figure 2E**.

### Differential kinetic behaviors of glioma cells at varied seeding densities

We employed GFP^+^ NPA high-grade glioma cells to investigate the impact of cell seeding density on oncostream formation (**Figure 3**). We utilized confocal time-lapse imaging to evaluate the role of initial cell density on oncostream formation in high-grade glioma cells at low (1 × 10^5 cells) and high (2 × 10^5 cells) densities over a 24-hour period (**Movies #1 (low density) and #2 (high density)**). At the start (0h) and after 24 hours (24h), time-lapse images and videos were acquired of distinct cell distributions. At low density, no oncostreams were observed, whereas high density conditions facilitated robust oncostream formation (**Figure 3A, J**). The TrackMate plugin in Fiji was used for tracking individual cell paths, revealing diverse patterns of cell movement over time (**Figure 3B, K**). Cell trajectories were refined using a Gaussian kernel filter in Julia, leading to smoother representations of cell movements (**Figure 3C, L**). Angular velocity distribution plots, showcasing the directional movement of cells, highlighted significant differences between the initial low- and high-density conditions (**Figure 3D, M**). Cell speed distributions and mean speed were calculated to understand the kinetic behavior relative to collective migration (**Figure 3E, N**). At a low seeding density, glioma cells were observed to move rapidly, averaging a speed of 34.76 µm/h. Despite this high speed, the cells did not exhibit a preferred direction of movement, nor did they demonstrate collective invasion patterns. In contrast, at the higher seeding density, cells showed a marked tendency to move collectively. Although they moved at a slower average speed of 11.57 µm/h, their movement was more directional and organized, indicative of a coordinated collective pattern, which is a characteristic behavior of oncostream formation (**Figure 3N**). Different collective migration patterns, categorized as streams, flocks, or swarms, were statistically analyzed. The analysis showed that low-density cells exhibited swarming behavior, while high-density cells predominantly formed streams (**Figure 3F, O**). Heatmaps visualized the velocity vector distributions, illustrating the dynamic contrast in oncostream formation under varying cell densities (**Figure 3G, P**). We further showed heatmap plots of the nematic correlation in relation to the proximity of neighboring cells, assessing uni- and bi-directional movements. These heatmaps revealed distinct spatial orientations and interaction dynamics at different densities (**Figure 3H, Q**). The frequency of cells relative to another cell’s position was quantified to map spatial distribution and interaction dynamics (**Figure 3I, R**). Collectively, these results demonstrate that seeding density significantly influences the kinetic behavior and collective migration patterns of aggressive glioma cells, with high density conditions favoring robust oncostream formation and organized collective invasion, contrasting the rapid but uncoordinated movement observed at low densities.

**Figure 3:**
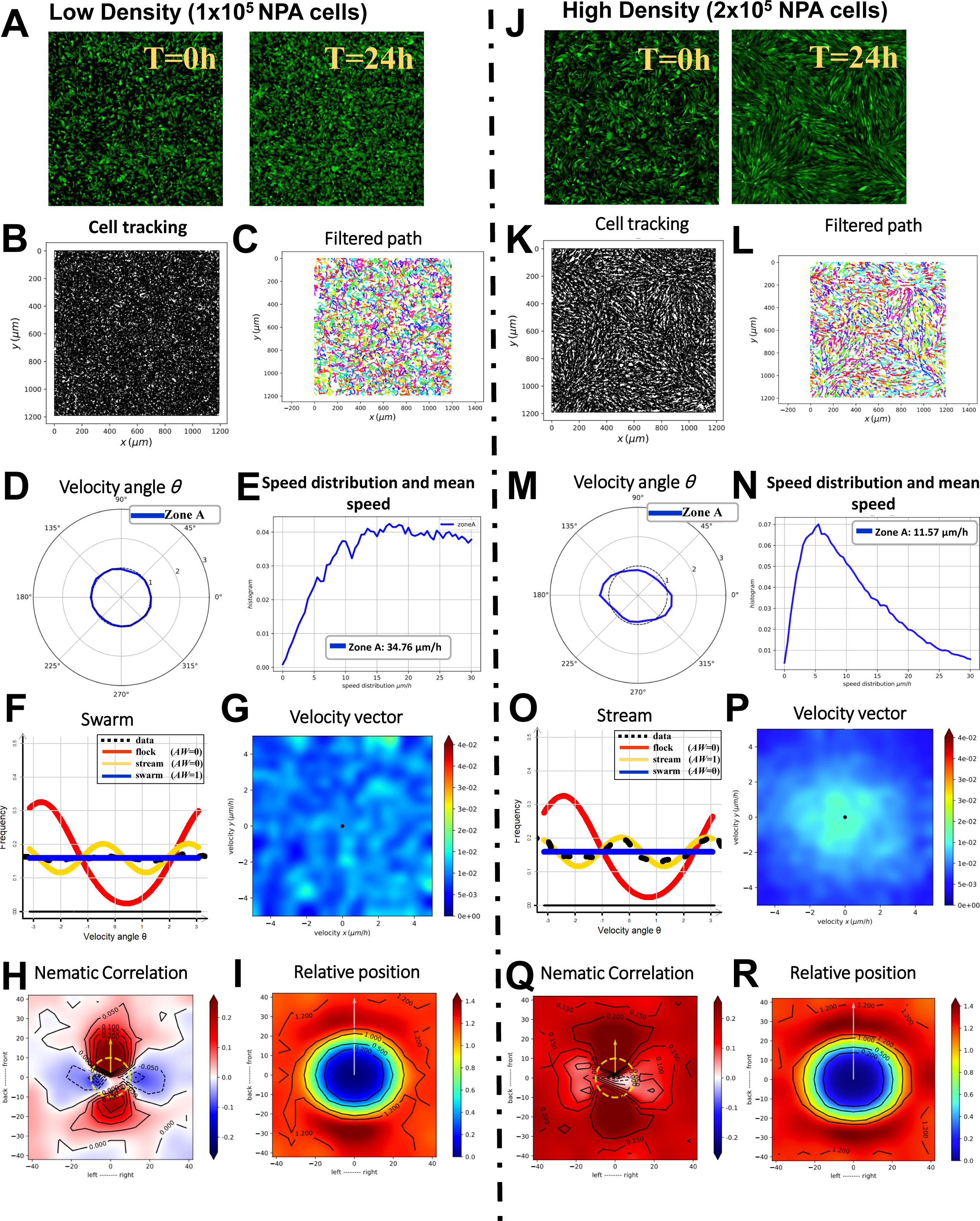
Influence of cell density on oncostreams formation in high-grade glioma cells. This figure demonstrates the effect of cell seeding density on the formation and dynamics of oncostreams. GFP+ NPA cells were seeded at two different densities, low (1 × 10^5 cells) and high (2 × 10^5 cells) and analyzed over a 24-hour period using confocal time-lapse imaging (see **Movies #1** and **#2**). **(A, J)** Time-lapse images captured at the start (0h) and after 24 hours (24h) illustrate the initial and subsequent cell distributions at low (no oncostreams) and high densities (full of oncostreams), respectively. **(B, K)** Using the TrackMate plugin in Fiji, the individual cell paths are tracked and displayed, elucidating the patterns of cell movement over time. **(C, L)** The cell trajectories are further refined using a Gaussian kernel filter (σ = 2, stencil of 9 points) in a Julia script, resulting in a smoothed representation of cell movement. **(D, M)** Angular velocity distribution plots present the directional movement of cells, with the angle θ indicating the tendency of cells to move in a specific direction for both low- and high-density conditions. **(E, N)** The distribution of cell speeds and the mean speed (μm/h) are calculated, providing insight into the overall kinetic behavior of the cells in the context of global collective migration. **(F, O)** Collective migration patterns are classified into streams (two peaks indicating two preferred angles of velocity), flocks (a single peak), or swarms (no preferred angle of velocity). Data distribution is estimated non-parametrically and analyzed for likelihood using different distribution models (ρ). The best fit model is selected based on the Akaike weight (AW). The frequency distributions reveal that low-density cells exhibit swarming behavior, whereas high-density cells tend to form streams. **(G, P)** Heatmaps visualize the velocity vector distributions throughout the entire time-lapse period, further contrasting the dynamics of oncostreams formation at varying cell densities. **(H, Q)** Heatmap plot of the nematic correlation relative to the proximity of neighboring cells are presented. A heat map plot of pairwise velocity correlation assesses the alignment of movement depending on the position of a nearby cell, (*xj)* in distinct spatial orientations (front-to-back and left-to-right). Positive correlation (+) indicates cells moving synchronously (red), while a negative correlation (-) suggests opposing directions (blue). **(I, R)** The frequency of cells in each zone relative to another cell’s position is quantified. From a cell’s reference point, we estimate the likelihood of another cell, (xj), being in proximity, mapping the spatial distribution and interaction dynamics between the cells at different densities.

### Spatiotemporal kinetics and collective behavior during oncostream formation

A time-lapse study over a 24-hour period revealed the stages of glioma cell organization into oncostreams (**Figure 4A**, **Movie #3).** Individual cell paths tracked across four-time intervals—0-6hr, 6-12hr, 12-18hr, and 18-24hr—highlighted dynamic migration patterns contributing to the formation of oncostreams (**Figure 4B**). Cell trajectories were refined using a Gaussian kernel filter, smoothing the cell paths and revealing a clearer progression of cellular organization (**Figure 4C**). The mean speed of cell movement decreased over time, indicative of a correlation with the transition from independent to collective behavior: 27.64 µm/h (0-6hr), 20.61 µm/h (6-12hr), 15.24 µm/h (12-18hr), and 11.69 µm/h (18-24hr) (**Figure 4D**). The observed decrease in cell speed correlates with increased cellular alignment and the formation of oncostreams, suggesting that the collective migration integral to oncostreams formation is a regulated process that evolves over time. Angular velocity distributions provided insights into the preferred direction of cell movement, showing a distinct transition over the observed periods (**Figure 4E**). Non-parametric estimates of data distribution, analyzed for the best model fit using Akaike weights, indicated a behavioral shift from swarming in early stages to stream formation indicative of organized oncostreams after 12 hours (**Figure 4F**). Further analysis of the dynamic interactions and spatial organization of glioma cells during oncostream formation globally is detailed in **Supplementary Figure S2**, which includes heatmaps of velocity distributions, pairwise correlations, nematic correlations, and spatial interaction frequency.

**Figure 4:**
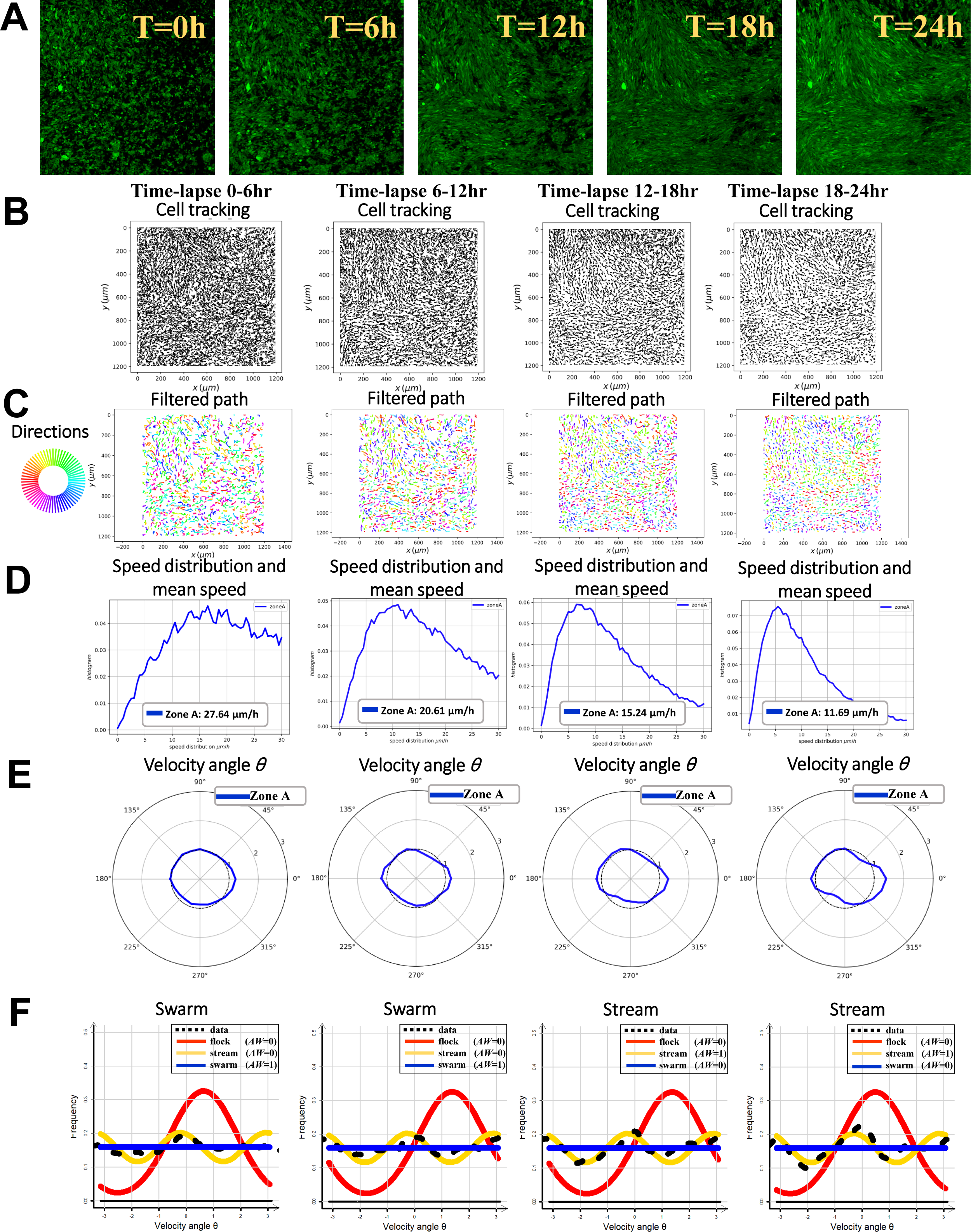
Spatiotemporal progression of oncostreams formation *in vitro*. This figure demonstrates the sequential self-formation of oncostreams at various developmental stages over 24hours (**see Movie #3**). **(A)** representative time-lapse images taken at the initial 0h, and at 6h, 12h, 18h and 24h intervals illustrate the development of cell organization into oncostreams. **(B)** The paths of individual cells are tracked across the time intervals (0-6hr, 6-12hr, 12-18hr, and 18-24hr), revealing the dynamic patterns of cell migration contributing to oncostream formation. **(C)** Cell trajectories are filtered using a Gaussian kernel filter (σ = 2, stencil of 9 points), resulting in a smoother representation of cell paths. **(D)** The distribution of cell speeds and the mean speed (μm/h) are calculated for each time interval, providing insight into the overall kinetic behavior of the cells during collective migration. **(E)** Angular velocity distribution, represented by the angle θ, demonstrates the preferred movement direction of cells during the specified periods. **(F)** Non-parametric estimates of data distribution are analyzed for model fit using different distribution types (ρ), with the Akaike weight (AW) determining the best-fitting model. Frequency distribution analysis indicates a transition from swarming behavior in the 0-6hr and 6-12hr periods to stream formation in cells after 12 hours.

In-depth localized analysis in distinct zones A, B, and C further elucidates the oncostream formation at different stages. This granular analysis, presented in Supplementary Figures S3-A to S3-E, demonstrates that the organization of oncostreams begins as early as the 0-6hr phase and continues to evolve through subsequent phases (6-12hr, 12-18hr, and 18-24hr). These supplementary figures reveal variations in cell movement speed, directionality, and spatial interactions within each specific zone, offering a comprehensive view of the structural evolution of oncostreams **(Supplementary Figures S3-A to S3-E)**.

This detailed spatiotemporal analysis captures the intricate process of oncostreams formation, where glioma cells initially exhibit a fast-paced, disorganized swarming pattern that gradually transitions to a slower, directed stream-like movement as they align into fascicle-like structures. The observed decrease in cell speed correlates with increased cellular alignment and the formation of oncostreams, suggesting that the collective migration integral to oncostreams formation is a regulated process that evolves over time.

### Collagenase-induced disruption of oncostreams and glioma cell migration dynamics

The incubation with 15 U/ml collagenase significantly disrupted pre-formed oncostreams, as demonstrated in a 15-hour time-lapse study (**Figure 5A**, **Movie #4**). GFP+ NPA glioma cells, after organizing into oncostreams over 36 hours, were exposed to sub-IC50 concentrations of collagenase, leading to the gradual disassembly of these structures (**Figure 5A**). A dose-response curve for collagenase confirmed that concentrations up to 100 U/ml did not reach half-maximal inhibitory concentration, indicating that the enzyme was effective at degrading the extracellular matrix without causing cell death (**Figure 5B**). Cell movement tracking post-treatment showed changes in migration patterns, transitioning from organized oncostreams to disorganized swarms due to ECM degradation (**Figure 5C**). Trajectories of individual cells were smoothed with a Gaussian kernel filter, facilitating the analysis of collective cell behavior following treatment (**Figure 5D**). Analysis of cell speed distributions and mean speeds over the treatment period indicated changes in migratory kinetics associated with structural dissociation of the oncostreams (**Figure 5E**). Angular velocity distribution analysis showed a loss of directional movement preference, suggesting a shift from coordinated migration to random, swarm-like behavior (**Figure 5F**). Likelihood analysis using different distribution models pointed to a transition from organized to disordered migration patterns during the collagenase treatment (**Figure 5G**). Velocity vector distribution heatmaps detailed the dynamics of oncostream dismantling across the entire observation period (**Figure 5H**). Pair-wise velocity correlations among neighboring cells were assessed, indicating alterations in the directionality of cell movement post-collagenase exposure (**Figure 5I**). A heatmap of nematic correlations relative to the proximity of neighboring cells depicted changes in directional alignment resulting from the collagenase treatment (**Figure 5J**). The frequency of relative positioning and spatial interactions between cell pairs over the treatment period was analyzed, mapping the response to the matrix-dependent dynamics post-collagenase exposure (**Figure 5K**). Altogether, these results indicate that collagenase treatment effectively dismantles the structural integrity of oncostreams by degrading the ECM, leading to a significant change in glioma cell migration from an organized, directional movement to a random and disorganized pattern. This suggests that the ECM plays a crucial role in maintaining the structure and collective invasion potential of oncostreams, providing insight into potential therapeutic strategies targeting the ECM.

**Figure 5:**
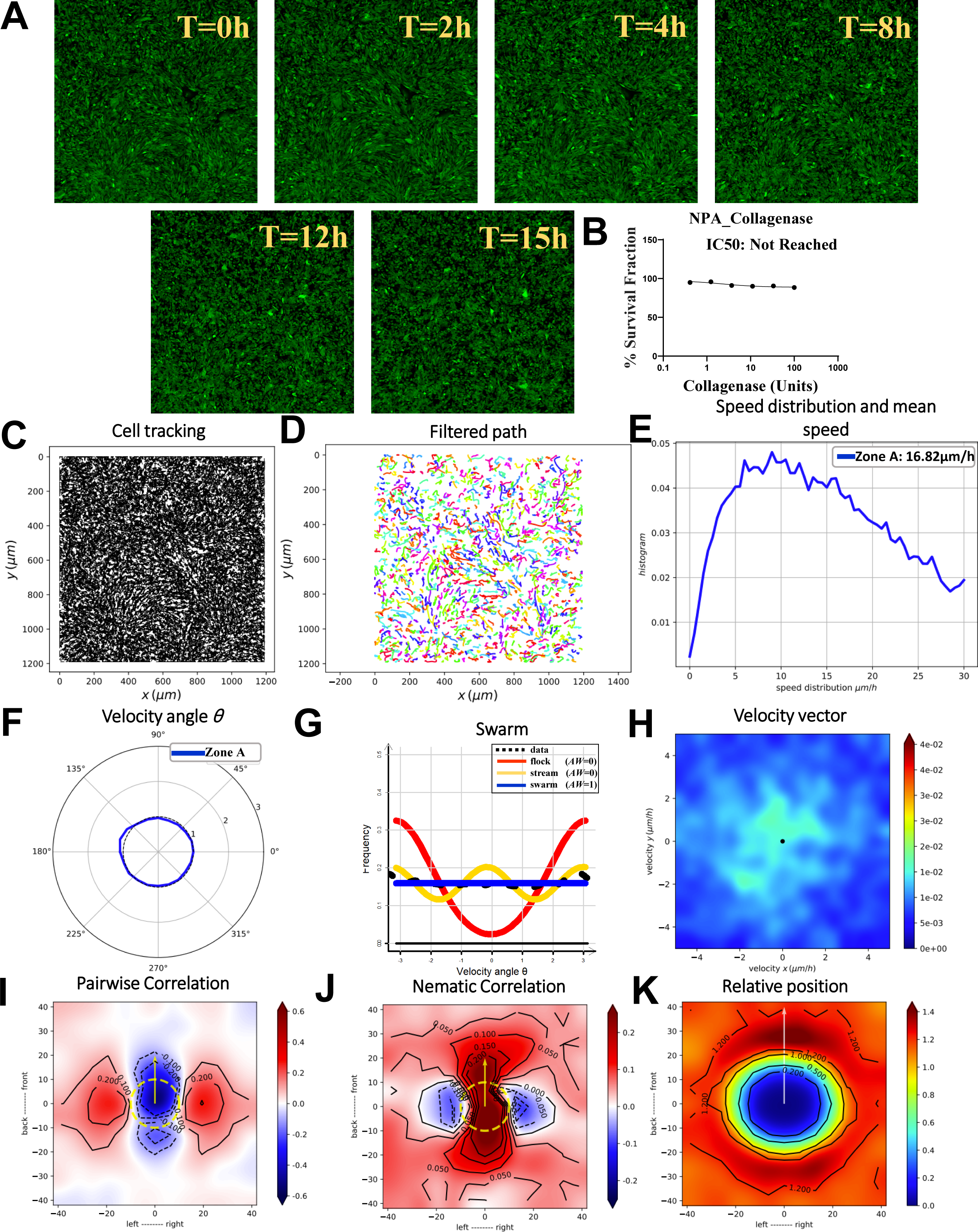
Collagenase dismantles oncostreams *in vitro* (15 U/ml). This figure illustrates the effect of 15 U/ml collagenase treatment on disrupting the structure of pre-formed aggressive and malignant oncostreams over a 15-hour period, as shown in **Movie #4**. **(A)** Following a 36-hour period post-seeding, where cells were allowed to self-organize into oncostreams, treatment was initiated with 15 U/ml collagenase (below the half-maximal inhibitory concentration, sub-IC50), intended to degrade the ECM without causing cell death. Representative time-lapse images captured at 0h, and subsequently at 2h, 4h, 8h, 12h, and 15h intervals, depict the gradual disassembly of oncostreams into disorganized swarms due to ECM degradation. **(B)** A dose-response curve is shown for GFP^+^ NPA glioma cells subjected to a three-fold serial dilution of collagenase ranging from 100 U/ml down to 0.411 U/ml. This range did not reach the IC50 value, suggesting that the concentration required for half-maximal inhibition is above 100 U/ml. **(C)** The paths of individual cells from 0 to 15 hours post-treatment are tracked to observe changes in migration patterns due to the presence of collagenase. **(D)** Cell trajectories are filtered using a Gaussian kernel filter (σ = 2, stencil of 9 points) for a smoother depiction of collective cell movement for further analysis. **(E)** Analysis of cell speed distributions and mean speeds (μm/h) over the 15-hour period provides insights into the migratory kinetics of cells undergoing structural dissociation. **(F)** The angular velocity distribution, quantified by angle θ, highlights the directional preferences of cell movement under the influence of collagenase. **(G)** Likelihood analysis of frequency distributions suggests a shift from coordinated oncostreams to disordered swarm behavior, characterized by a lack of preferred direction, over a 15-hour collagenase treatment period. **(H)** Heatmaps demonstrate the velocity vector distributions for the complete time-lapse duration (0-15hr), further illustrating the dynamic dismantling of oncostreams. **(I)** Pair-wise velocity correlations among neighboring cells are assessed, considering the relative positions (**xi - xj**) and calculating the direction correlation (**ωi to ωj**), with axes representing the cell positions and distances in µm. The dotted yellow line indicates the average size of the centered cell. **(J)** A heatmap of nematic correlations related to the proximity of neighboring cells provides a visual representation of directional alignment as affected by collagenase treatment. **(K)** The frequency of relative positioning between pairs of cells across each time-lapse segment is analyzed in response to collagenase, estimating the probability of cell proximity and mapping spatial interactions and matrix-dependent dynamics post-treatment.

### Impact of TC-I-15 on integrin-mediated glioma cell adhesion and oncostream development

The integrin antagonist TC-I-15 significantly impeded the assembly of glioma cell structures, inhibiting oncostream formation as captured over a 20-hour period (**Figure 6A**, **Movie #5)**. TC-I-15 was used at 5 µM after a 6-hour cell adhesion period, effectively blocking integrin interactions without causing cell death, as evidenced by time-lapse confocal imaging (**Figure 6A**). A dose-response curve was established for GFP+ NPA glioma cells with a two-fold serial dilution of TC-I-15, showing effective inhibition below the half-maximal inhibitory concentration (**Figure 6B**). Cell migration paths after TC-I-15 treatment revealed altered migratory patterns, transitioning from organized to disorganized behavior (**Figure 6C**). Application of a Gaussian kernel filter smoothed cell trajectories, providing a clear depiction of changes in collective movement (**Figure 6D**). Analysis of cell speed distributions and mean speeds revealed alterations in migration kinetics, highlighting cytoskeletal changes in response to TC-I-15 (**Figure 6E**). Angular velocity distribution, quantified by angle θ, indicated the absence of preferred directional cell movement after treatment (**Figure 6F**). Frequency distribution analysis comparing treated cells with controls suggested a shift from a coordinated to a disorganized swarm behavior over the treatment period (**Figure 6G**). Heatmaps of velocity vector distributions throughout the time-lapse demonstrated the dynamic effect of TC-I-15 on oncostreams disruption (**Figure 6H**). Pair-wise velocity correlations between cells were quantified, showing a decrease in coordinated movement directionality (**Figure 6I**). Heatmaps of nematic correlations relative to neighboring cell proximity revealed changes in cellular alignment due to TC-I-15 (**Figure 6J**). The frequency of cells in close proximity within each time-lapse frame was calculated, mapping the spatial distribution and interaction dynamics altered by TC-I-15 (**Figure 6K**). TC-I-15 effectively disrupts α2β1 integrin-mediated adhesion, leading to the inhibition of oncostreams formation and altering the collective migration dynamics of glioma cells. Moreover, these results provides insights into the therapeutic potential of TC-I-15 in modulating the invasive behavior of glioma cells by targeting integrin-mediated interactions.

**Figure 6:**
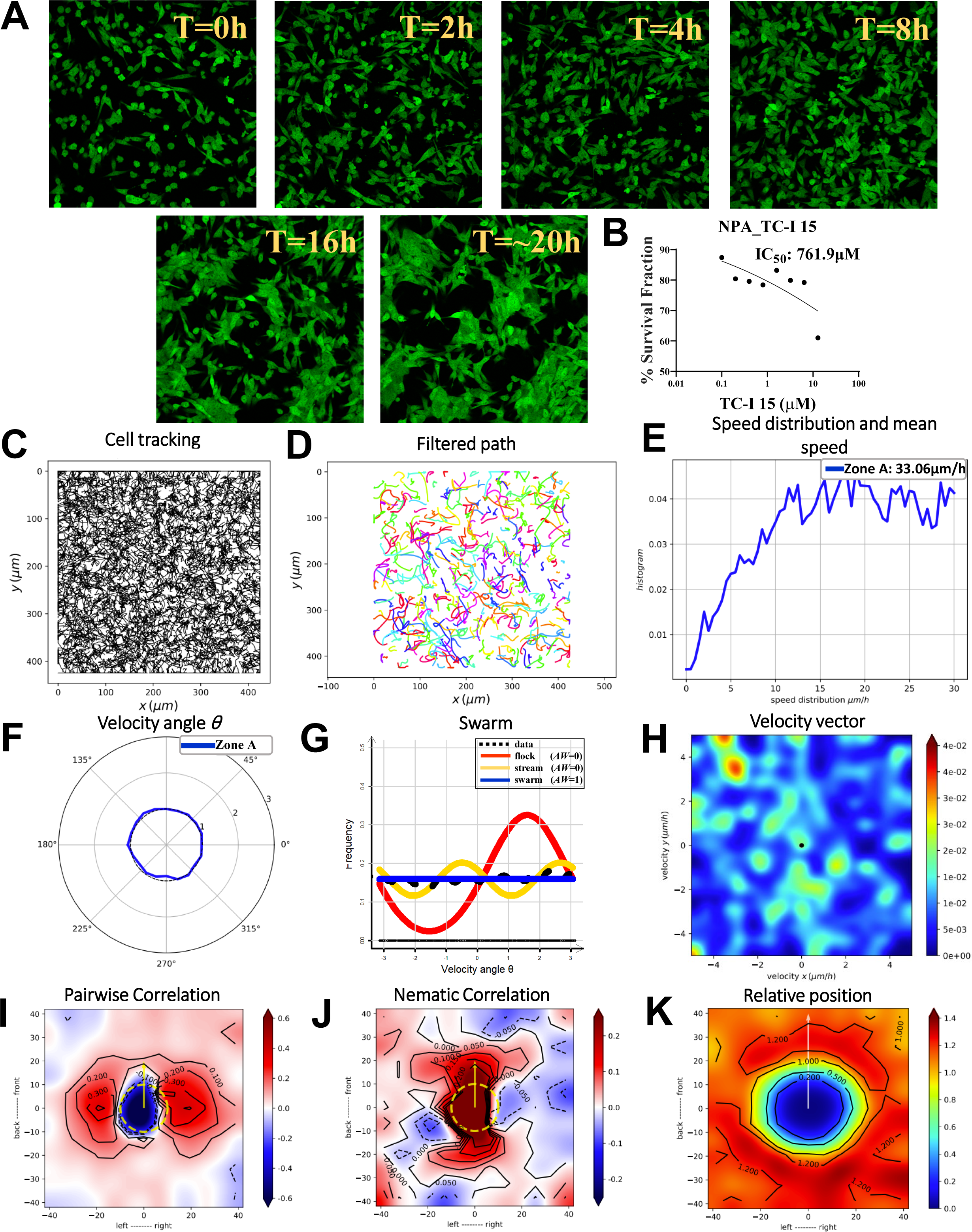
TC-I-15 inhibits α2β1 integrin-mediated adhesion and oncostream formation. This figure explains the effect of the integrin antagonist TC-I-15 on glioma cell adhesion and oncostream formation. Time-lapse movies were recorded over 20 hours, as shown in **Movie #5**. **(A)** Cells were allowed to adhere for 6 hours post-seeding before the introduction of 5 µM TC-I-15 (sub-IC50). This was intended to block integrin interactions without inducing cell death. Time-lapse images taken at 0h, and at 2h, 4h, 8h, 16h, and 20h intervals, show the inhibited assembly of cell structures (oncostreams), resulting in disorganized swarms due to the disruption of integrin-mediated adhesion. **(B)** A dose-response curve for GFP^+^ NPA glioma cells is shown, following a two-fold serial dilution of TC-I-15 from 12.8μM to 0.1μM. **(C)** Cell migration paths from 0 to 20 hours post-treatment are tracked to reveal alterations in migratory patterns resulting from the integrin antagonist intervention. **(D)** A Gaussian kernel filter (σ = 2, stencil of 9 points) is applied to the cell trajectories to obtain a smoothed/filtered representation of collective movement. **(E)** Cell speed distributions and mean speeds (μm/h) are analyzed over the 20-hour period to determine the migration kinetics of cells as they undergo cytoskeletal structural changes. **(F)** Angular velocity distribution, indicated by angle θ, is used to understand the preferred directions of cell movement in response to the integrin antagonist. **(G)** Frequency distribution analysis, performed to compare treated cells with controls, suggests a transition from a coordinated pattern to a disorganized swarm behavior lacking a preferred direction after 20 hours of TC-I-15 treatment. **(H)** Heatmaps of velocity vector distributions for the entire time-lapse period (0-20hr) highlight the disruption of oncostream development due to integrin antagonist presence. **(I)** Pair-wise velocity correlations between neighboring cells, considering their relative positions (xi - xj), are quantified to determine the correlation of directional velocities (ωi and ωj). Axes mark the cell positions and distances in µm, with a dotted yellow line representing the average size of a centered cell. **(J)** Heatmaps display the nematic correlation relative to the proximity of neighboring cells, showing TC-I-15 treatment alters cellular alignment. **(K)** The frequency of cells in close proximity within each time-lapse frame is calculated in response to TC-I-15, mapping the spatial distribution and interaction dynamics of the cells.

### Cytochalasin D disruption of actin polymerization and oncostreams organization

Cytochalasin D, known to impede actin polymerization, was observed to disrupt the structural organization of oncostreams in GFP^+^ NPA glioma cells (**Figure 7A**, **Movie #6)**. Upon the administration of 5 µM Cytochalasin D, time-lapse imaging over a 1-hour period demonstrated rapid cellular responses, transitioning from organized streams to disorganized swarms (**Figure 7A**). A dose-response curve established that the IC50 value for Cytochalasin D was 15.43 µM. The used concentration of 5 µM effectively avoided cell death while impairing cell behavior and oncostream integrity (**Figure 7B**). Cell migration paths were tracked before and after Cytochalasin D treatment, revealing a marked alteration in migratory patterns, indicative of the inhibitor’s impact on glioma dynamics (**Figure 7C**). Trajectories were refined using a Gaussian kernel filter, elucidating the collective movements as cells underwent structural disorganization (**Figure 7D**). Angular velocity distribution analysis showed a lack of preferred direction of cell movement post-treatment (**Figure 7E**). A significant decrease in cell motility was noted post-Cytochalasin D treatment, with average speeds reduced to 5.8 µm/h, emphasizing the inhibitor’s effect on cellular dynamics (**Figure 7F**). Heatmaps of velocity vector distributions illustrated the dynamic disassembly of oncostreams under the influence of the treatment over the hour (**Figure 7G**). Likelihood analysis suggested a transition from the coordinated movement characteristic of oncostreams to a disorganized swarm behavior with no apparent preferred direction after 1 hour of Cytochalasin D treatment (**Figure 7H**). Pair-wise velocity correlations among neighboring cells showed changes in directional velocities, indicating a disruption of cellular alignment (**Figure 7I**). Nematic correlation heatmaps provided a visual representation of how collective cellular alignment was affected by Cytochalasin D (**Figure 7J**). The spatial interactions and the likelihood of cell proximity were recalculated post-treatment, mapping the actin-dependent dynamics in response to Cytochalasin D (**Figure 7K**). These results suggest that the inhibitory effect of Cytochalasin D on actin polymerization leads to the dismantling of the intricate oncostream structures, thereby corroborating the critical role of the actin cytoskeleton in maintaining the collective invasion mechanism of glioma cells. Furthermore, these findings enhance our understanding of the cytoskeletal dependencies in oncostream integrity and highlight potential therapeutic strategies targeting the structural components during glioma cell invasion.

**Figure 7:**
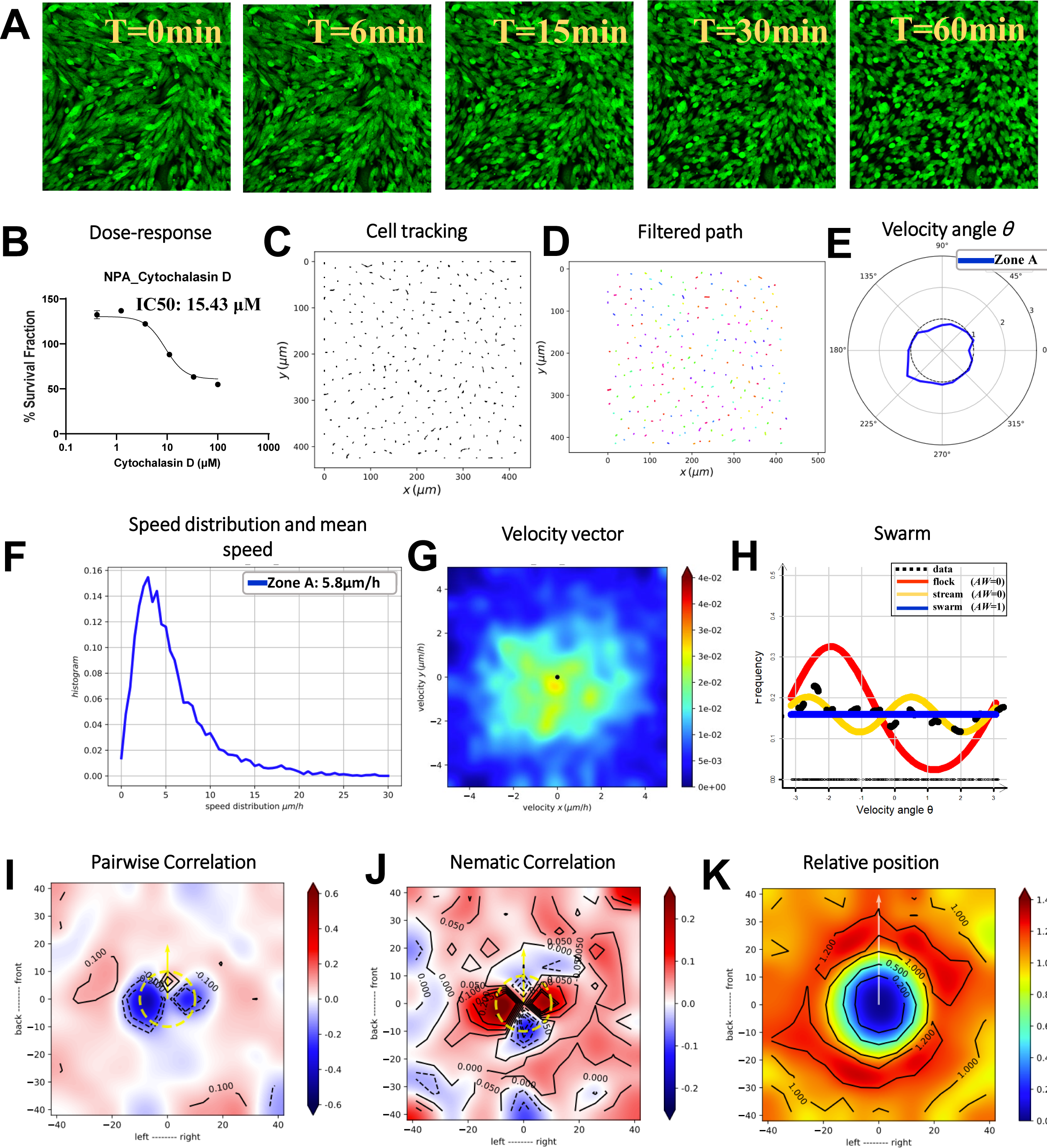
Impact of the actin cytoskeleton in oncostreams formation using Cytochalasin D. This figure explains the effect of Cytochalasin D on disrupting oncostream formation by inhibiting actin polymerization in GFP^+^ NPA glioma cells. Cytochalasin D, a cell-permeable inhibitor known to block actin polymerization, was administered at a concentration of 5 µM. Time-lapse imaging was conducted over a 1-hour period to capture the immediate cellular response, with frames acquired at 1.5-minute intervals. The resulting time-lapses are compiled in **Movie #6**, demonstrating the rapid and dynamic cellular changes during this period. **(A) S**equential images at 0, 6, 15, 30, and 60 minutes post-Cytochalasin D treatment show a distinct disruption in the organization of the oncostreams, with cells shifting from an organized stream to a disorganized swarm. **(B)** A dose-response curve for GFP^+^ NPA glioma cells was plotted following a three-fold serial dilution of Cytochalasin D, ranging from 100 µM to 0.411 µM. The IC50 value was determined to be 15.43 µM. For the treatment demonstrates in the time-lapse movie, a sub-IC50 concentration of 5 µM was used to avoid cell death while still observing the effects on cell behavior and oncostream integrity. **(C)** Tracking of cell migration paths from 0 to 1-hour post-treatment reveals alterations in migratory patterns due to the disruption of actin polymerization. **(D)** Cell trajectories are smoothed using a Gaussian kernel filter (σ = 2, stencil of 9 points), providing a refined representation of collective cell movements. **(E)** Angular velocity distribution, represented by angle θ, is used to understand the preferred directions of cell movement post-Cytochalasin D treatment. **(F)** Post-Cytochalasin D treatment, there was a noticeable decrease in cell movement, with the average speed dropping to 5.8 µm/h, highlighting the compound’s impact on cell motility and the dismantling of oncostreams. **(G)** Heatmaps of velocity vector distributions throughout the entire 1-hour time-lapse showcase the dynamic disassembly of oncostreams under treatment. **(H)** Likelihood analysis of frequency distributions indicates a shift from coordinated oncostreams to disorganized swarm behavior with no preferred direction after 1-hour of Cytochalasin D treatment. **(I)** Pair-wise velocity correlations among neighboring cells are assessed by determining their relative positions (**xi - xj**) and calculating the direction correlation (**ωi to ωj**). Axes represent the cell positions and distances in µm, with a dotted yellow line indicating the average size of the centered cell. **(J)** A heatmap of nematic correlations, based on the proximity of neighboring cells, represents how collective cellular alignment is affected by Cytochalasin D treatment. **(K)** The frequency of relative positioning between pairs of cells throughout each time-lapse interval is analyzed, estimating the likelihood of proximity, and mapping spatial interactions as well as actin-dependent dynamics in response to Cytochalasin D treatment.

### Myosin II inhibition by p-nitro Blebbistatin disassembles oncostreams

P-nitro Blebbistatin at a 5 µM concentration significantly disrupted the organization of oncostreams in GFP^+^ NPA glioma cells, as depicted in the transition from organized streams to disorganized swarms over a 1-hour period following treatment (**Figure 8A**, **Movie #7)**. This disruption highlights the critical role of myosin II in maintaining oncostream integrity (**Figure 8A**). Conversely, treatment with 4-HAP did not dismantle the oncostream structure, suggesting a differential role or efficacy in the inhibition mechanisms between the two compounds **(Supplementary Figure S4)**. The IC50 value for p-nitro Blebbistatin was determined to be 307.3 µM, and a sub-IC50 concentration of 5 µM was utilized to minimize cytotoxicity while impairing myosin II activity (**Figure 8B**). Tracking of cell migration paths post-treatment with p-nitro Blebbistatin displayed significant changes in migratory patterns (**Figure 8C**), while similar tracking under 4-HAP treatment did not indicate such drastic changes, as cell motility remained relatively unchanged **(Supplementary Figure S4E)**. Application of a Gaussian kernel filter to the cell trajectories resulted in a smoother depiction of cell movements, aiding the analysis of changes in collective behavior (**Figure 8D**). Angular velocity distribution analysis after p-nitro Blebbistatin treatment supported the loss of directed cell movement (**Figure 8E**), a feature that was not observed in the 4-HAP treated cells, which showed no significant alteration in directional movement **(Supplementary Figure S4F)**. A marked reduction in cell motility was quantified post-p-nitro Blebbistatin treatment, with average cell speed reduced to 7.53 µm/h (**Figure 8F**), whereas 4-HAP treatment did not result in a significant change in cell speed **(Supplementary Figure S4E)**. Heatmaps illustrating velocity vector distributions captured the rapid disassembly of oncostreams under the influence of p-nitro Blebbistatin (**Figure 8G**), whereas 4-HAP treatment-maintained velocity vector distributions indicative of intact oncostreams **(Supplementary Figure S4H)**. Likelihood analysis of frequency distributions post-p-nitro Blebbistatin treatment indicated a transition from coordinated to disorganized behavior (**Figure 8H**), a phenomenon not observed with 4-HAP treatment **(Supplementary Figure S4G)**. Assessment of pairwise velocity correlations and nematic correlations heatmaps showed a decrease in directional alignment among neighboring cells post-p-nitro Blebbistatin treatment (**Figures 8I and 8J**), contrasting with the sustained alignment in the 4-HAP treated cells **(Supplementary Figure S4I and S4J)**. Finally, the spatial distribution and interaction dynamics between cells were quantified, illustrating the treatment’s impact on myosin-dependent dynamics (**Figure 8K**), while 4-HAP treatment did not show significant impact on these dynamics **(Supplementary Figure S4K)**. Collectively, these findings illustrate that myosin II inhibition by p-nitro Blebbistatin effectively dismantles oncostreams in glioma cells, highlighting the pivotal role of non-muscle myosin II ATPases (IIA and IIB) in organized collective cell movement. The lack of similar effects with 4-HAP treatment suggests specificity in the myosin II inhibitory mechanisms and highlights the potential of selective targeting in therapeutic intervention.

**Figure 8:**
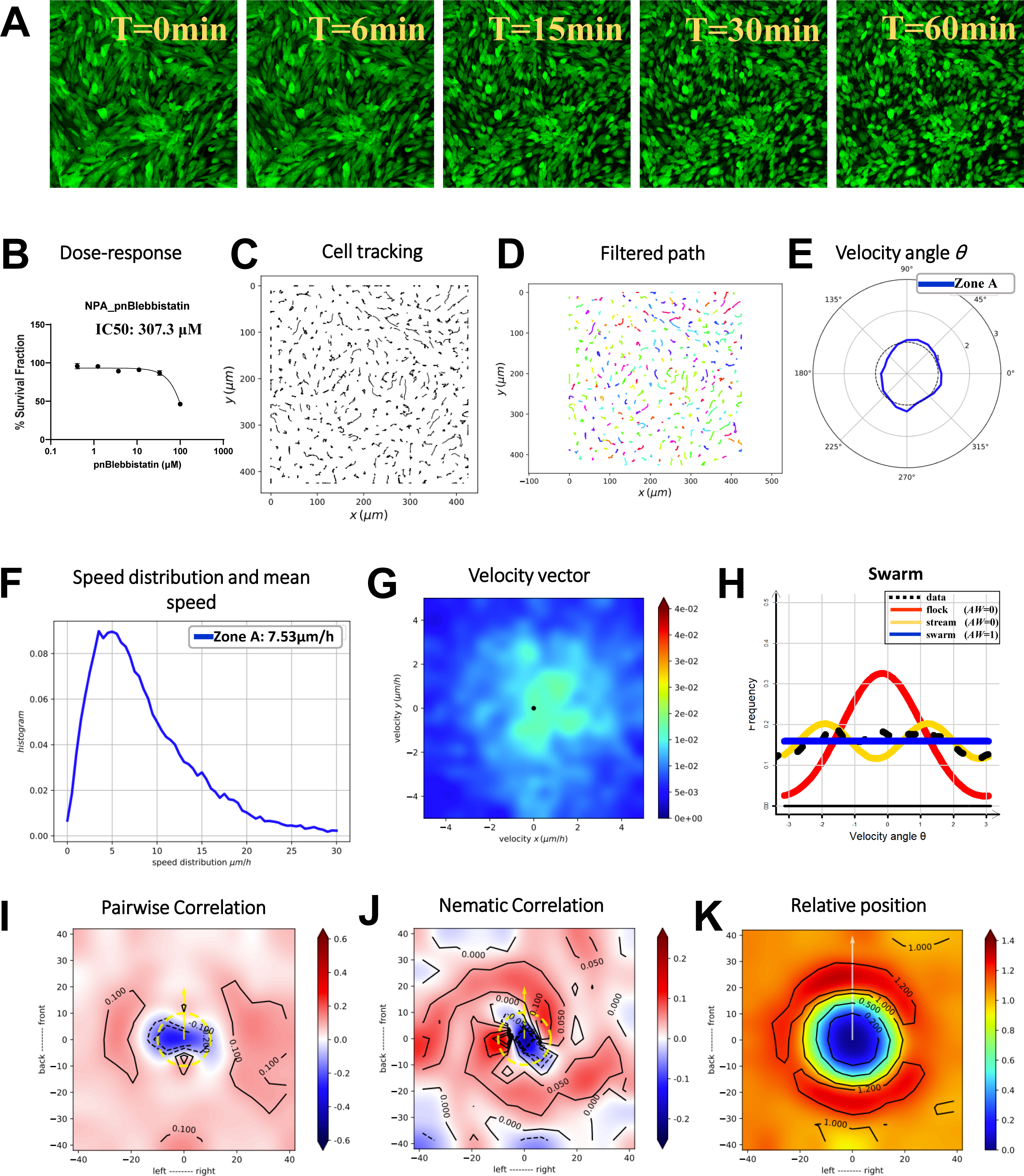
Role of myosin II in oncostreams formation using p-nitro Blebbistatin. This figure presents the effect of p-nitro Blebbistatin on oncostream formation by specifically inhibiting myosin II activity in GFP^+^ NPA glioma cells. P-nitro Blebbistatin, a cell-permeable inhibitor of myosin II ATPase activity, was administered at a concentration of 5 µM. Time-lapse imaging over a 1-hour period captured the immediate cellular response, with frames acquired at 1.5-minute intervals, highlighting the inhibitor’s rapid effect. The corresponding time-lapses are compiled in **Movie #7**, showcasing the dynamic changes induced by myosin II inhibition. **(A)** Time-lapse images at 0, 6, 15, 30, and 60 minutes after p-nitro Blebbistatin treatment reveal a notable disruption in oncostream organization. The cells are observed transitioning from an ordered stream to a disorganized swarm, implicating myosin II as a critical player in maintaining oncostream structure. **(B)** A dose-response curve was plotted for NPA glioma cells subjected to a three-fold serial dilution of p-nitro Blebbistatin, ranging from 100 µM down to 0.411 µM. The IC50 value was determined to be 307.3 µM. For the treatment, a concentration of 5 µM below the IC50 was chosen to avoid cell death while still disrupting myosin II activity. **(C)** The migration paths of individual cells from 0 to 1-hour post-treatment were tracked, displaying alterations in migratory patterns due to the inhibition of myosin II. **(D)** The cell trajectories were processed using a Gaussian kernel filter (σ = 2, stencil of 9 points), resulting in a smoothed representation of cell movements. **(E)** The angular velocity distribution, represented by angle θ, was analyzed to understand changes in the preferred directions of cell movement post-p-nitro Blebbistatin treatment. **(F)** There was a significant reduction in cell motility following p-nitro Blebbistatin application, with the average speed decreasing to 7.53 µm/h, emphasizing the inhibitor’s role in cellular movement and oncostream disruption. **(G)** Heatmaps of velocity vector distributions across the 1-hour time-lapse period clearly depict the dynamic disassembly of oncostreams under the influence of p-nitro Blebbistatin. **(H)** Likelihood analysis of frequency distribution indicates a shift from coordinated oncostreams to disorganized swarm behavior with no preferred direction post-pnBB treatment. **(I)** Pair-wise velocity correlations among neighboring cells were assessed, considering their relative positions (xi - xj) and calculating the directional correlation (ωi to ωj). The axes represent the cell positions and distances in µm, with a dotted yellow line indicating the average size of the centered cell. **(J)** Heatmaps of nematic correlations based on the proximity of neighboring cells, represents how collective cellular alignment is altered by p-nitro Blebbistatin treatment. **(K)** The frequency of relative positioning between cells across each time-lapse frame was calculated, estimating the likelihood of proximity, and mapping spatial interactions and myosin-dependent dynamics in response to p-nitro Blebbistatin.

### Calcium chelation by BAPTA-AM alters oncostream formation and cell motility

Time-lapse confocal imaging over a 16-hour period delineated the impact of BAPTA-AM on oncostream formation in GFP+ NPA glioma cells (**Figure 9A-J**, **Movie #8** for untreated and **Movie #9** for treated cells). In untreated conditions, cells organized into oncostreams, while BAPTA-AM treatment resulted in the absence of these structures, with cells displaying disorganized swarming behavior (**Figure 9A, J**). The TrackMate plugin in Fiji tracked individual cell paths, showing that BAPTA-AM altered cell trajectories significantly (**Figure 9B, K**). Trajectories were smoothed using a Gaussian kernel filter, which provided a clear contrast in cell movement between untreated and treated cells (**Figure 9C, L**). Angular velocity distribution plots indicated a clear directional movement in untreated cells, which was disrupted by BAPTA-AM treatment, reflecting a loss of collective migration (**Figure 9D, M**). A significant reduction in cell motility was observed following BAPTA-AM treatment, with the average speed dropping from 8.04 µm/h in untreated cells to 4.58 µm/h in treated cells (**Figure 9E, N**, and **Supplementary Figure S5**). Non-parametric data distribution analysis using Akaike weight (AW) demonstrated that untreated cells-maintained stream-like structures, whereas BAPTA-AM-treated cells exhibited swarming behavior (**Figure 9F, O**). Heatmaps of velocity vector distributions throughout the time-lapse period showcased the contrasting dynamics between untreated and treated groups, highlighting the disassembly of oncostreams upon BAPTA-AM treatment (**Figure 9G, P**). Nematic correlation heatmaps related to the proximity of neighboring cells showed that untreated cells moved synchronously, while BAPTA-AM treatment disrupted this alignment, leading to asynchronous movement (**Figure 9H, Q**). The frequency of relative cell positioning was quantified, revealing a spatial distribution and interaction dynamics shift in response to BAPTA-AM (**Figure 9I, R**). A dose-response curve for BAPTA-AM determined that the chosen concentration of 5 µM for treatment did not reach the IC50 value, ensuring cell viability while affecting cellular behavior (**Figure 9S**). Our result further reveals that calcium chelation by BAPTA-AM significantly alters oncostream formation and cell motility in glioma cells, disrupting organized collective migration and leading to disorganized swarming behavior, thus highlighting the crucial role of calcium signaling in the regulation of oncostream dynamics and cellular movement.

**Figure 9:**
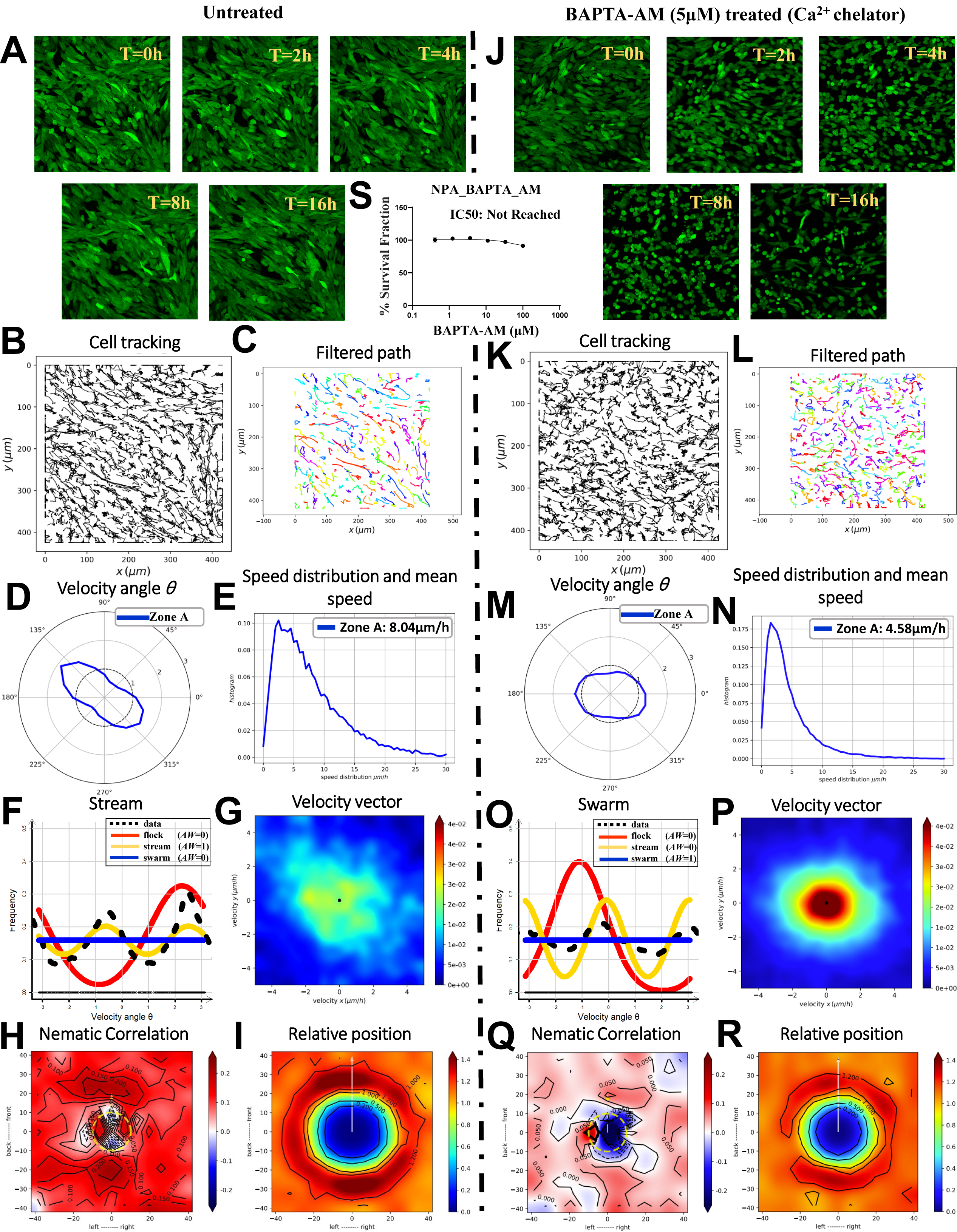
Effect of BAPTA-AM treatment on oncostream. This figure demonstrates the formation and dynamics of oncostreams with and without BAPTA-AM treatment. The cells were monitored over a 16-hour period using confocal time-lapse imaging (refer to **Movie #8** for untreated and **Movie #9** for BAPTA-AM treated). **(A, J)** Time-lapse images at 0, 2, 4, 8, and 16 hours compare the cell organization between untreated and BAPTA-AM-treated cells, demonstrating the presence or absence of oncostreams, respectively. **(B, K)** Individual cell paths were tracked using the TrackMate plugin in Fiji, elucidating the movement patterns over time and the influence of BAPTA-AM treatment on these trajectories. **(C, L)** The cell trajectories were further filtered using a Gaussian kernel filter (σ = 2, stencil of 9 points) using Julia-based script, displaying movement for both untreated and BAPTA-AM-treated groups. **(D, M)** Angular velocity distribution plots present the directional movement of cells, with the angle θ revealing the tendency of cells to move in a specific direction in both untreated and BAPTA-AM-treated conditions. **(E, N)** The distribution of cell speeds and the mean speed (μm/h) are calculated, there was significant reduction cell motility following BAPTA-AM treatment, with the average speed decreasing to 4.58 µm/h from 8.04 µm/h under untreated conditions. **(F, O)** Data distributions are estimated non-parametrically and analyzed for the best fit using Akaike weight (AW). Frequency distributions suggest that BAPTA-AM-treated cells exhibit swarming behavior, while untreated cells are more likely to retain stream-like structures. **(G, P)** Heatmaps show the velocity vector distributions throughout the entire time-lapse period, highlighting the contrasting dynamics of oncostreams formation in untreated versus BAPTA-AM-treated cells. **(H, Q)** Heatmap plots of nematic correlation relative to the proximity of neighboring cells assess the alignment of movement. Positive correlation (+, shown in red) indicates cells moving synchronously, while negative correlation (-, shown in blue) indicates cells moving in opposite directions. **(I, R)** The frequency for each zone of the relative position between two cells is quantified. Estimations of the probability of having another cell (xj) in proximity are made from the perspective of one cell, mapping the spatial distribution and interaction dynamics influenced by BAPTA-AM treatment. **(S)** A dose-response curve was plotted for NPA glioma cells subjected to a three-fold serial dilution of BAPTA-AM, ranging from 100 µM down to 0.411 µM. For the treatment, a concentration of 5 µM was chosen for treatment based on the observed dose-response curve, even though the IC50 value was not reached.

### Neurotransmitter modulation of oncostream dynamics: effects of glutamate and histamine

Confocal time-lapse imaging over a 16-hour period delineated the impact of glutamate and histamine on the formation and dynamics of oncostreams (**Figure 10A, J**; **Movie #10** for glutamate and **Movie #11** for histamine treatment). In the presence of glutamate and histamine, cells displayed distinct organization patterns, with glutamate increasing cell motility and histamine reducing it compared to control conditions (**Figure 10A, J**). Using the TrackMate plugin in Fiji, individual cell paths were tracked, revealing the influence of glutamate and histamine on movement patterns (**Figure 10B, K**). Cell trajectories were refined with a Gaussian kernel filter, clarifying the movement patterns after neurotransmitter treatments (**Figure 10C, L**). Angular velocity distribution plots indicated the preferred movement directions, with variations observed between glutamate and histamine treatments (**Figure 10D, M**). Cell motility analysis showed an increase to 11.74 µm/h after glutamate treatment and a decrease to 6.88 µm/h post-histamine treatment, in contrast to the 8.04 µm/h observed in untreated cells (**Figure 10E, N**, and **Supplementary Figure S5**). Non-parametric estimation and analysis using Akaike weight (AW) highlighted that both glutamate and histamine treatments preserved the stream-like structures, with differing effects on cell motility (**Figure 10F, O**). Heatmaps of velocity vector distributions showcased the varied oncostream dynamics between cells treated with glutamate and histamine (**Figure 10G, P**). Nematic correlation heatmaps provided insights into the alignment of movement, with differing patterns of synchronization in response to each neurotransmitter (**Figure 10H, Q**). The spatial interactions and probability distributions of cell proximity were quantified, mapping the effects of glutamate and histamine treatments on cell distribution and interaction dynamics (**Figure 10I, R**). Dose-response curves for NPA glioma cells to glutamate and histamine treatments were plotted, confirming that IC50 values were not reached. Selected concentrations of 1 mM glutamate and 5 µM histamine were used based on these curves to avoid toxicity (**Figure 10S, T**). These results demonstrates that neurotransmitter modulation, specifically through glutamate and histamine, distinctly affects oncostream dynamics in glioma cells, with glutamate enhancing and histamine reducing cell motility, thereby providing valuable insights into the complex interplay between neurotransmitter signaling and the structural and motility aspects of oncostream formation. Both neurotransmitters allowed the preservation of stream-like structures, indicating that oncostream formation can be influenced by neurotransmitter signaling without complete disruption of collective cell organization.

**Figure 10:**
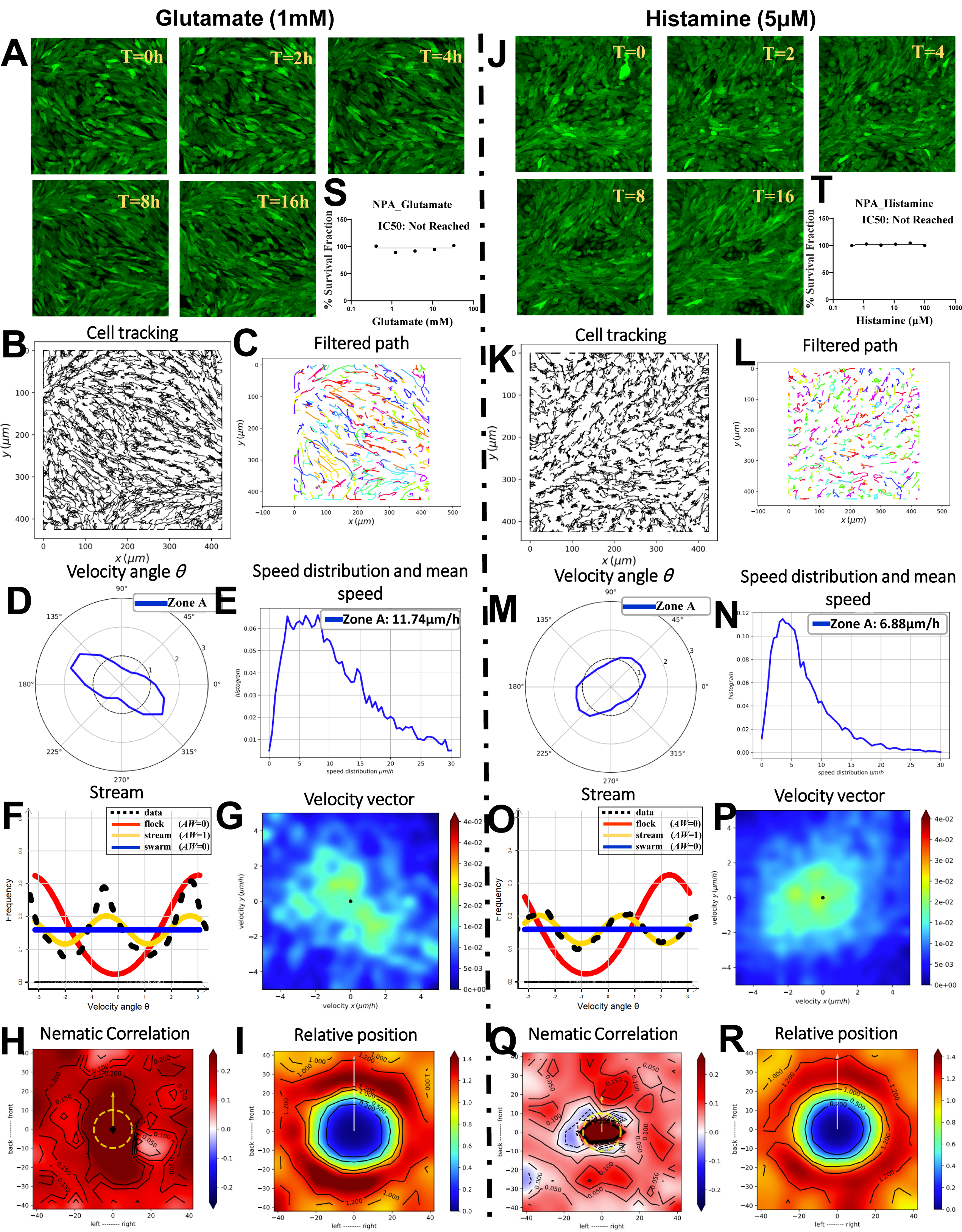
Effect of neurotransmitters glutamate and histamine on oncostreams. This figure demonstrates the formation and dynamics of oncostreams in the presence of glutamate and histamine treatment, monitored over a 16-hour period using confocal time-lapse imaging (refer to **Movie #10** for glutamate and **Movie #11** for histamine treatment). **(A, J)** Comparative time-lapse images at 0, 2, 4, 8, and 16 hours display cell organization in cells treated with glutamate and histamine, highlighting dynamics behavior of oncostreams. **(B, K)** Individual cell paths were tracked using the TrackMate plugin in Fiji, elucidating movement patterns over time and the influence of both treatments on these trajectories. **(C, L)** Cell trajectories were further filtered using a Gaussian kernel filter (σ = 2, stencil of 9 points) using Julia-based script, displaying movement patterns in treated groups. **(D, M)** Angular velocity distribution illustrate directional movement, with the angle θ indicating cells’ preferred movement directions under each treatment. **(E, N)** The distribution of cell speeds and the mean speed (μm/h) are analyzed, revealing a significant increase in cell motility to 11.74 µm/h following glutamate treatment, while a decrease to 6.88 µm/h was observed following histamine treatment, compared to 8.04 µm/h under untreated conditions. **(F, O)** Non-parametric estimation and analysis using Akaike weight of data distributions show that both treatment groups maintain stream-like structures, albeit with variations in cell motility. **(G, P)** Heatmaps display velocity vector distributions over the entire time-lapse period, contrasting oncostream dynamics in cells treated with glutamate versus histamine. **(H, Q)** Nematic correlation heatmaps assess alignment of movement, with red indicating positive (synchronous) correlation and blue indicating negative (opposite direction) correlation. **(I, R)** The spatial distribution and interaction dynamics influenced by the treatments are quantified (I, R), mapping frequency and probability distributions of cells in proximity. **(S, T)** A dose-response curves for NPA glioma cells were plotted for glutamate (ranging from 100 mM to 0.411 mM) and histamine (ranging from 100 µM to 0.411 µM). IC50 values were not reached for either treatment. Concentrations of 1 mM glutamate and 5 µM histamine were chosen based on these curves, ensuring effective treatment without reaching toxic levels.

### Modulation of oncostream dynamics by Rho GTPase signaling

In exploring the regulatory mechanisms of oncostream formation, we assessed the role of Rho GTPase signaling. Treatment with Rho-Activator I led to notable alterations in oncostream integrity. Comparative analysis of time-lapse images revealed a significant reorganization of cell structures, transitioning from well-formed streams to more disorganized patterns upon activator treatment **(Supplementary Figure S6A, K).** This was further substantiated by cell trajectory analysis, showing marked changes in cell movement **(Supplementary Figure S6B, L),** and quantified by a significant decrease in cell motility, with average cell speeds reduced after activator treatment **(Supplementary Figure S6D, N).** Conversely, the application of a Rho-Inhibitor had a distinct impact on oncostream dynamics. Time-lapse imaging captured a less pronounced but observable effect on cell organization **(Supplementary Figure S7A),** with trajectory smoothing highlighting subtler changes in movement patterns **(Supplementary Figure S7C).** The analysis of angular velocity distribution and cell speed further demonstrated the inhibitory effects on cell motility, albeit to a lower extent than the activator **(Supplementary Figure S7D, E).** Both treatments affected the cell motility, as evidenced by speed distribution **(Supplementary Figures S6D, N and S7E),** suggesting that Rho GTPase activity is important for the maintenance of oncostream motility and invasion. Moreover, analysis of pairwise velocity correlations and nematic correlations after Rho-Inhibitor treatment provided insights into the collective movement dynamics, reflecting alterations in cell alignment and interaction **(Supplementary Figures S6H, I, R, S and S7H, I).** These findings highlight the complexity of Rho GTPase signaling in oncostream motility and maintenance. While Rho-Activator I disrupts oncostream integrity and cell motility significantly **(Supplementary Figure S6),** Rho-Inhibitor also impacts these dynamics but in a distinct manner, hinting at the delicate balance of activation and inhibition within this signaling pathway **(Supplementary Figure S7).** Such insights pave the way for targeted therapeutic strategies that modulate the Rho GTPase activity in cancer progression.

## Discussion

In our previous studies, we described the existence of oncostreams, dynamic multicellular structures that play a pivotal role in the intra-tumoral distribution of cells and potentially facilitate the collective invasion of the normal brain. These oncostreams, characterized by a unique molecular signature with a prominent overexpression of COL1A1, were directly correlated with glioma aggressiveness.(*26, 27, 38*)

Herein, we have taken a step further by modeling glioma oncostreams *in vitro*. This innovative approach allowed us to explore the cellular dynamics of glioma oncostream behavior, within an optimized environment to evaluate the impact of various pharmacological agents on oncostream formation and organization.

Our findings highlight the significance of cell density and morphology in predicting self-organization in gliomas. This observation aligns with our previous work, where we demonstrated that high-grade gliomas, characterized by their dense oncostreams, displayed a more aggressive phenotype.(*26, 38*) Furthermore, our *in vitro* experiments highlight the role of the extracellular matrix, particularly collagen, in facilitating oncostream formation. We observed a complete dismantling of oncostreams in response to collagenase treatment. Our *in vitro* data are consistent with our previous *in vivo* observations where COL1A1 was found to be overexpressed in oncostreams, and whose down regulation eliminated oncostreams from Sleeping Beauty induced tumors *in vivo*.(*26*)

The in-depth statistical analyses using *in-house* developed scripts for Julia programming and R Studio has decoded complex migratory patterns, delimiting glioma cells into distinct behaviors such as flocks, streams, and swarms. This classification not only corroborates our earlier work (*26*) but also provides a quantifiable framework to assess the efficacy of potential therapeutic interventions aimed at disrupting these patterns.

The testing of pharmacologic agents in our models has uncovered new layers of oncostream regulation. Agents such as Cytochalasin D and p-nitro Blebbistatin illuminated the essential involvement of the cytoskeleton, with actin polymerization and myosin II activity emerging as critical components for maintaining oncostream integrity. Complementing this, the significance of calcium signaling in cell motility and oncostream structure was highlighted by our use of calcium chelators like BAPTA-AM. This aligns with earlier findings that Ca^2+^ signals are pivotal in controlling cell migration through their regulation of forward movement and cell adhesion.(*39-42*) Moreover, the modulation of cell behavior by neurotransmitters such as glutamate and histamine demonstrated the influence of biochemical signaling on tumor cell dynamics. Significantly, TC-I-15, an integrin antagonist, disrupts α2β1 integrin-mediated adhesion in glioma cells, inhibiting oncostream formation and modifying collective cell migration, a crucial factor in glioma progression, thus underscoring its therapeutic potential against glioma invasiveness. Targeting multiple integrins can enhance the radiosensitivity of collectively invading cancer cells, overcoming their resistance to radiotherapy and DNA damage.(*43*)

It has been previously demonstrated that the migration patterns of cancer cells at the center of tumors are spatially organized and driven by actin cytoskeleton.(*44*) The invasive phenotype of glioma cells is governed by a complex interplay of factors, signaling cascades, and interactions between cellular and ECM components.(*45-50*) This complexity presents challenges in targeting glioma invasion.(*45, 51*) Our data sheds light on this intricate regulation through oncostream-focused *in vitro* modeling and pharmacological testing, demonstrating the multifaceted nature of glioma cell migration and the influence of cellular components and ECM on it.

Given the persistent challenge in improving GBM patient outcomes, our findings emphasize the challenges of current surgical techniques, which are often limited by the diffuse nature of glioma invasion.(*2, 9, 11, 14, 20-22*) Our in vitro model, therefore, stands as a promising platform for the discovery of novel therapeutic agents. By simulating the invasion process through oncostreams, we can scrutinize the effects of drugs on glioma dissemination. The *in vitro* disruption of oncostreams by agents targeting the ECM, cytoskeletal components, and calcium signaling pathways highlights the potential of these targets in impeding the invasive spread of glioma cells. Our findings attest to the critical role of intercellular communication in oncostream integrity and suggest that modulation of these factors can halt or reverse tumor progression.

In summary, our *in vitro* oncostream model will serve as a valuable tool for advancing our understanding of glioma invasion and testing therapeutic strategies aimed at undermining the cellular and molecular foundations of GBM malignancy. The insights gained from this study hold the promise of guiding future research towards interventions that can effectively disrupt the collective invasion and oncostreams, thereby offering hope for more effective GBM therapies.

## Conclusions

In summary, our study offers a novel perspective on glioma invasion by successfully modeling oncostreams *in vitro*, shedding light on the intricate spatiotemporal collective dynamics of glioma cells. Our thorough investigation highlights the impact of various pharmacological agents on oncostream formation, maintenance, and behavior. The significance of targeting oncostreams and their associated molecular pathways emerges as a promising strategy to inhibit glioma progression and combat the aggressive nature of glioblastoma. Significantly, our *in vitro* model highlights the importance of targeting oncostreams and their molecular pathways, not only as a strategy to curb glioma progression and invasion into the normal brain parenchyma but also in the broader context of combating metastasis in various cancers. Our study transcends traditional scratch and migration assays by offering quantitative analysis of biophysical factors like velocity, speed, and likelihood, thereby enhancing our comprehension of cancer invasion and metastasis. These insights pave the way for novel anti-invasive and anti-metastatic therapies, further highlighting the potential for developing more effective treatments across a spectrum of invasive and metastatic tumors. Our model thus contributes crucially to the advancement of cancer therapy and holds promise for significantly improving patient outcomes and survival rates in glioma and other metastatic cancers.

## Supporting information

Movies related to the manuscript

## Acknowledgments

We would like to acknowledge the funding support provided by [grant/funding information]. This work was supported by National Institutes of Health/National Institute of Neurological Disorders and Stroke (NIH/NINDS) grants: R37-NS094804, R01-NS105556, R01-NS122536, R01-NS124167, R01-NS122165, and R21-NS123879-01 to M.G.C.; NIH/NINDS grants: R01-NS076991, R01-NS096756, R01-NS082311, R01-NS122234, R01-NS127378 to P.R.L.; National Institute of Biomedical Imaging and Bioengineering (NIH/NIBI): R01-EB022563, National Cancer Institute (NIH/NCI) U01-CA224160, RNA Biomedicine grant: F046166 to M.G.C., and Rogel Cancer Center at The University of Michigan G023089 to M.G.C. NIH/National Cancer Institute grant R01-CA243916 to PRL; Ian’s Friends Foundation grant G024230, Leah’s Happy Hearts Foundation grant G013908, Pediatric Brain Tumor Foundation grant G023387, Smiles for Sophie Forever Foundation and ChadTough Foundation grant G023419 to M.G.C. and P.R.L. We are also grateful to Karin Muraszko and Aditya Pandey for outstanding academic leadership, Melissa for excellent administrative assistance and Marta Dzaman for superb technical and valuable input throughout this study.

## Author contributions

SMF wrote the first draft of this manuscript, with overall guidance and revisions of MGC and PRL. SMF acquired and analyzed the confocal data. SMF prepared the figures with input from MGC and PRL. The manuscript was reviewed and edited collaboratively by SMF and PRL. JEC, BS, MLV, AC, and GA provided assistance to SMF in performing the experiments. PRL contributed the H&E for the upper panel of Figure 1A and 1B. SM reviewed the mathematical and statistical analysis conducted by SMF. All authors participated in the work and approved the final version of the manuscript for submission.

## Data Availability Statement

The supplementary movies and their corresponding descriptions, which are integral to our study are available for public access. These materials can be found at the following digital repository: Zenodo (https://doi.org/10.5281/zenodo.10381916). Additionally, the complete Julia and R Studio scripts and codes used for the statistical analysis of oncostreams in our study are also publicly accessible. These resources are crucial for understanding the computational aspect of our research and can be accessed at: Zenodo (https://doi.org/10.5281/zenodo.7587235).

## Conflict of Interest Statement

The authors declare no conflicts of interest related to this work.

## Notes

https://doi.org/10.5281/zenodo.10381916

https://doi.org/10.5281/zenodo.7587235

### Competing Interest Statement

The authors have declared no competing interest.

