## Supplemental Data for "Modeling Glioma Oncostreams In Vitro: Spatiotemporal Dynamics of their Formation, Stability, and Disassembly"

**Running title:** Collective Behavior in Glioblastoma: Unraveling the Dynamics of Brain Cancer Invasion

Figure S1

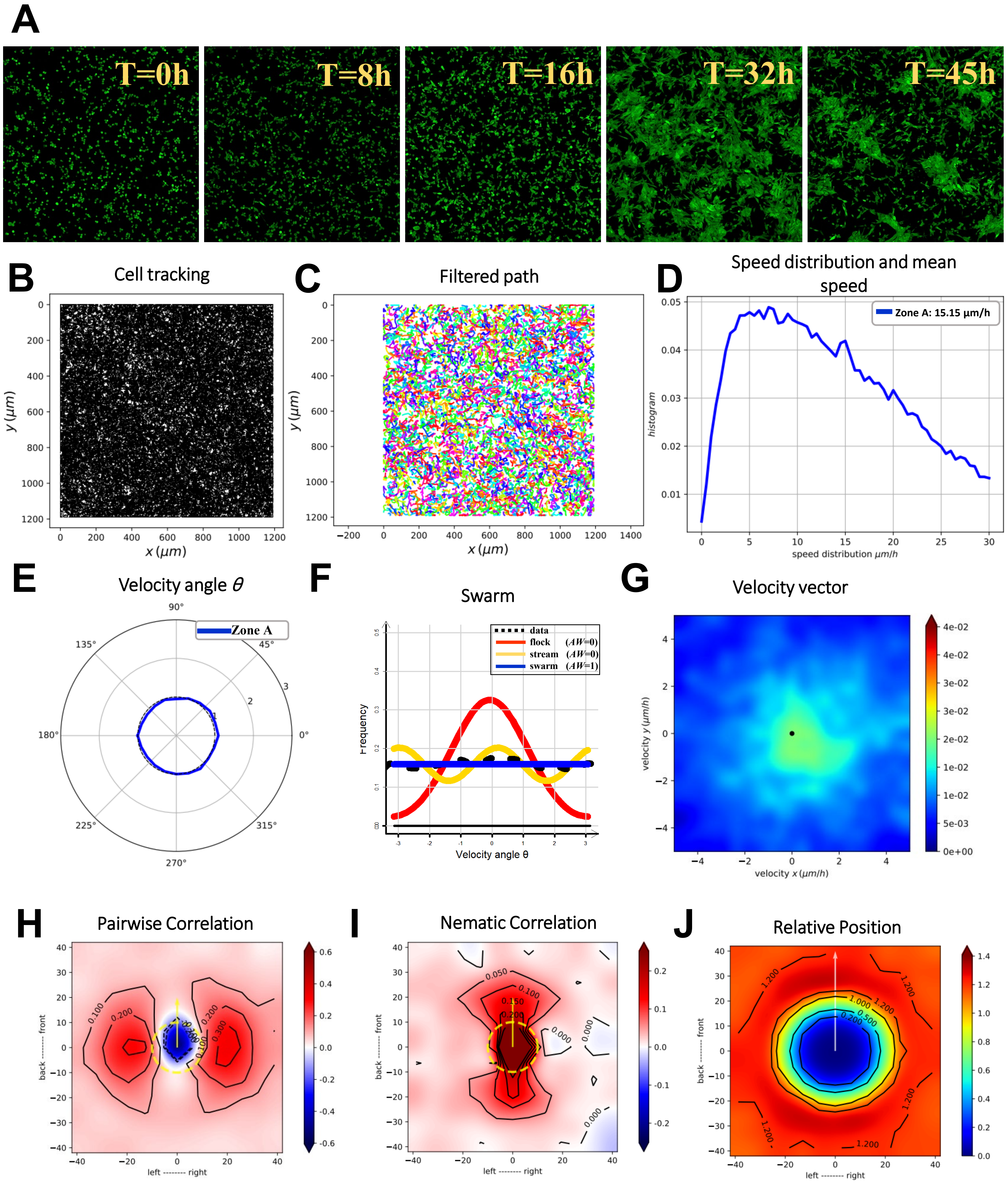

A
T=0h
T=8h
T=16h
T=32h
T=45h
B
Cell tracking
y (μm)
x (μm)
C
Filtered path
y (μm)
x (μm)
D
Speed distribution and mean speed
Zone A: 15.15 μm/h
histogram
speed distribution μm/h
E
Velocity angle θ
Zone A
90°
135°
45°
180°
0°
225°
315°
270°
F
Swarm
data
flock (AW=0)
stream (AW=0)
swarm (AW=1)
Frequency
Velocity angle θ
G
Velocity vector
velocity y (μm/h)
velocity x (μm/h)
H
Pairwise Correlation
back -------- front
left -------- right
I
Nematic Correlation
back -------- front
left -------- right
J
Relative Position
back -------- front
left -------- right

**Supplementary Files**

#### Supplementary Figures Legends

**Figure_S1: Lack of oncostream formation on poly-d-lysine coated culture dishes without laminin.** This figure illustrates the inability of $GFP^{+}$ NPA glioma cells to organize into oncostreams when cultured on poly-D-lysine coated glass-bottom dishes in the absence of laminin. Over a period of 45 hours, time-lapse imaging was performed, and images were taken at 10-minute intervals to document the behavior of glioma cell migration and organization. The corresponding time-lapses are compiled in **Movie #12**, showcasing the dynamic oncostream organization compromised without laminin. **(A)** Initial and subsequent images at 0, 8, 16, 32, and 45 hours show cells forming dense clusters rather than structured oncostreams, highlighting the essential role of laminin in oncostream architecture. **(B)** Analysis of individual cell migration paths from the start to the end of the observation period reveals a disorganized, swarm-like pattern of movement attributed to the lack of a laminin substrate. **(C)** Cell trajectories were smoothed using a Gaussian kernel filter ($\sigma = 2$, stencil of 9 points), offering a clearer view of the migration paths. **(D)** Quantification of migration speed shows a reduced average velocity of 15.15 μm/h, indicating a decrease in cell motility on poly-D-lysine alone. **(E)** The angular velocity distribution is examined to discern any preferred directionality in cell movement, with angle $\theta$ serving as the measurement parameter. **(F)** Frequency distribution analysis via likelihood estimation underscores the random, disoriented swarm-like behavior of the cell, lacking a preferred direction. **(G)** Velocity vector distribution heatmaps from the 45-hour time-lapse reveal the extent and nature of glioma cell clustering in response to the absence of laminin coating. **(H)** Velocity correlations between neighboring cells are investigated, considering their relative positions and measuring directional correlation to elucidate the influence of cell-to-cell interactions in the observed pattern. **(I)** Nematic correlation heatmaps based on cell proximity display the collective orientation, or lack thereof, reinforcing the conclusion that laminin is crucial for the alignment and formation of oncostreams. **(J)** The frequency of relative cell positioning is calculated for each time frame, providing insight into the likelihood of cell interaction and spatial organization, which is compromised without laminin.

### Figure S2

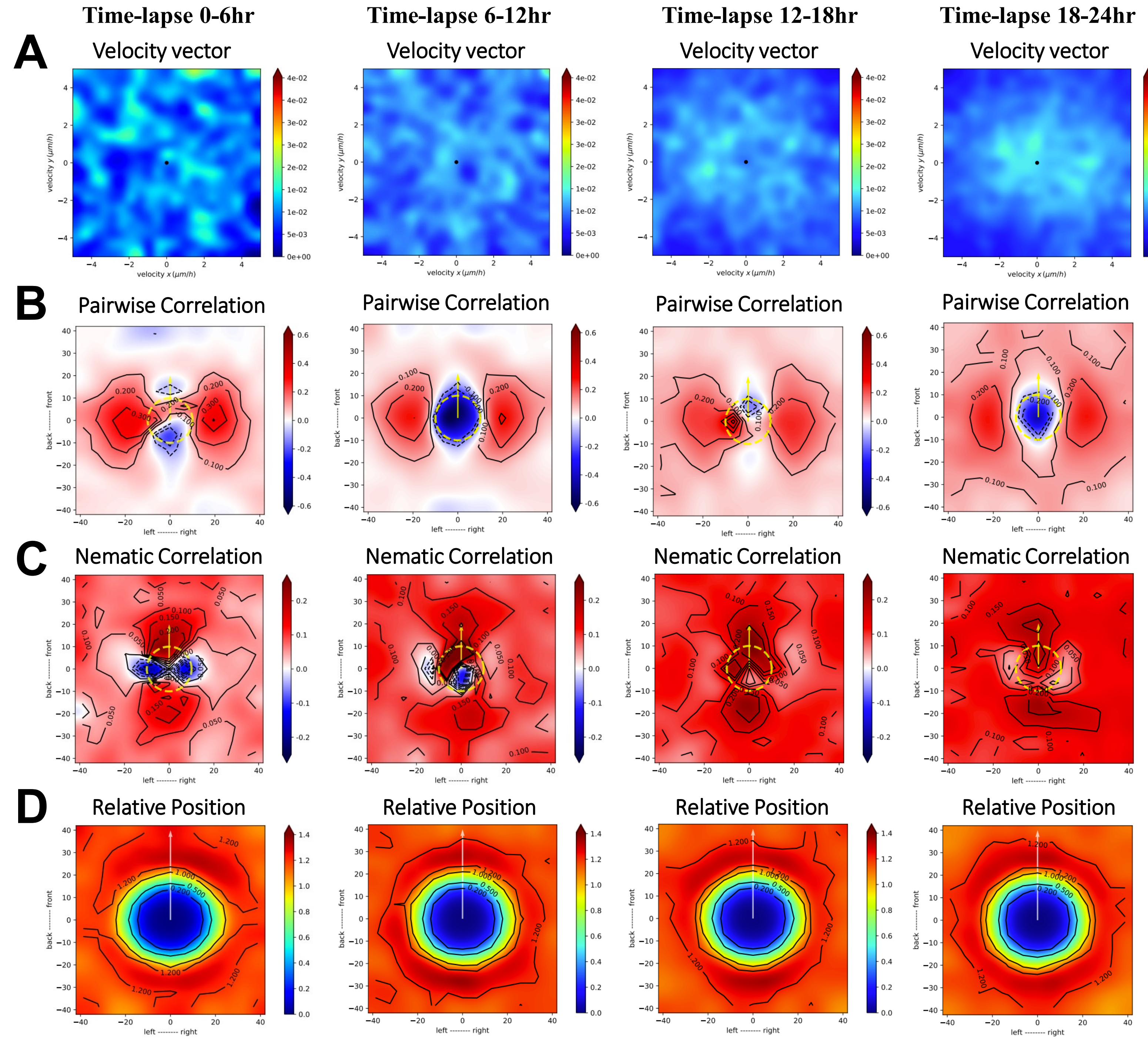

Time-lapse 0-6hr
Time-lapse 6-12hr
Time-lapse 12-18hr
Time-lapse 18-24hr
A
Velocity vector
velocity y (μm/h)
velocity x (μm/h)
B
Pairwise Correlation
back -------- front
left -------- right
C
Nematic Correlation
D
Relative Position

#### Supplementary Files

**Figure_S2: Extended analysis of oncostream formation and migration dynamics over 24 hours. (A)** Heatmaps of velocity vector distributions for the entire duration of the time-lapse (0-24hr) provide a comprehensive view of the dynamic oncostream formation. These visualizations allow for the observation of changes in movement speed and directionality at various stages: early (0-6hr), mid (6-12hr), late (12-18hr), and final (18-24hr). **(B)** Pair-wise velocity correlations between neighboring cells are quantified, taking into account their relative positions (xi - xj) and directional correlation (ωi to ωj) over the same time intervals. The axes denote cell positions and distances in micrometers, with a dotted yellow line marking the average size of a central cell, illustrating the coordination in cell motility during oncostream development. **(C)** Heatmaps of nematic correlations based on the proximity of adjacent cells highlight the alignment and directionality of cell movements across the different phases of oncostream formation (0-6hr, 6-12hr, 12-18hr, and 18-24hr), providing insights into the collective behavior of cells. **(D)** The frequency of relative positioning between cell pairs for each segment of the time-lapse is evaluated, calculating the likelihood of proximity to map spatial interactions and matrix-dependent dynamics throughout the observed intervals, thereby elucidating the structural evolution of oncostreams.

Figure S3-A

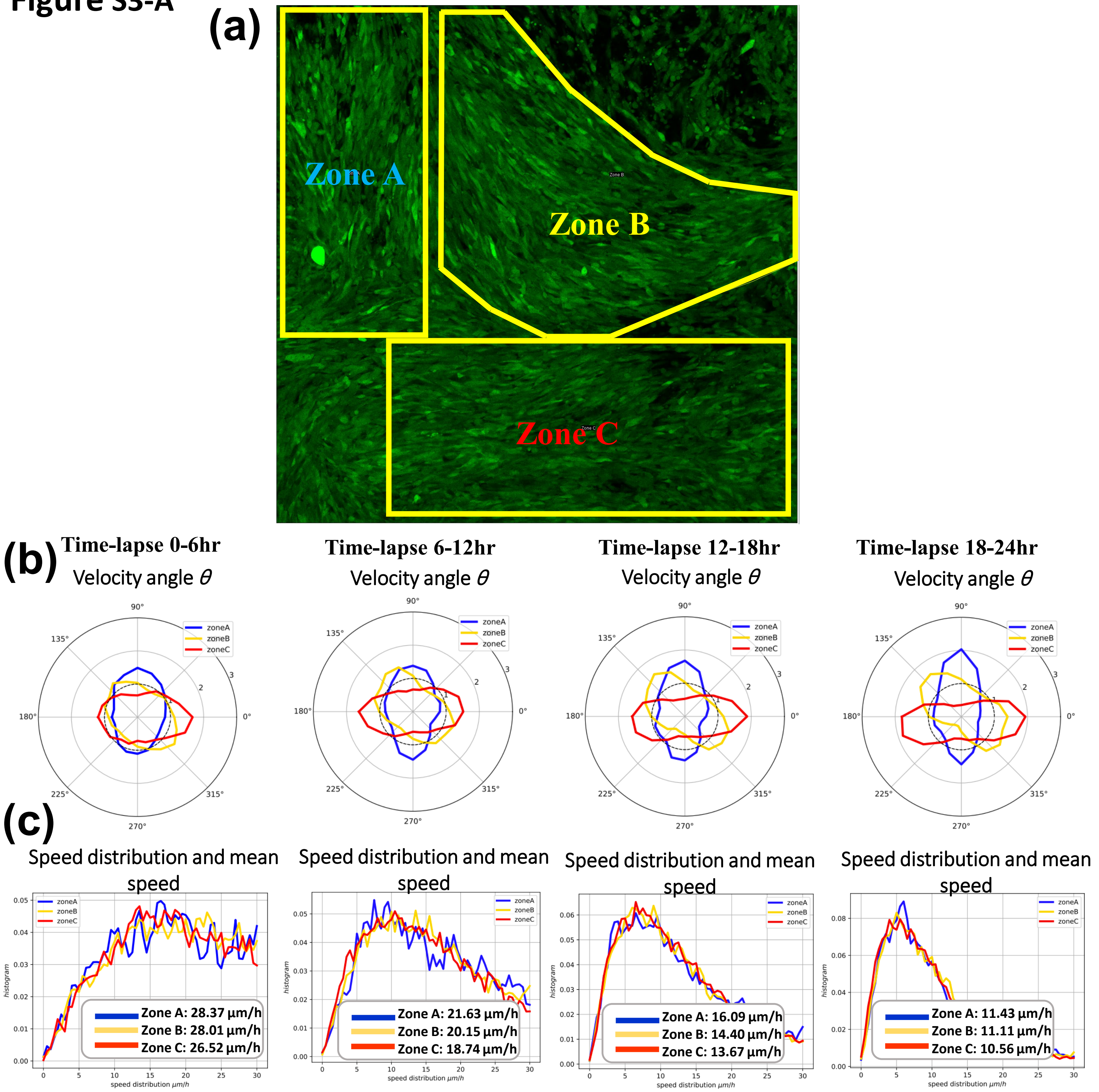
(a)
Zone A
Zone B
Zone C
(b)
Time-lapse 0-6hr
Velocity angle θ
Time-lapse 6-12hr
Velocity angle θ
Time-lapse 12-18hr
Velocity angle θ
Time-lapse 18-24hr
Velocity angle θ
zoneA
zoneB
zoneC
(c)
Speed distribution and mean speed
Zone A: 28.37 μm/h
Zone B: 28.01 μm/h
Zone C: 26.52 μm/h
Speed distribution and mean speed
Zone A: 21.63 μm/h
Zone B: 20.15 μm/h
Zone C: 18.74 μm/h
Speed distribution and mean speed
Zone A: 16.09 μm/h
Zone B: 14.40 μm/h
Zone C: 13.67 μm/h
Speed distribution and mean speed
Zone A: 11.43 μm/h
Zone B: 11.11 μm/h
Zone C: 10.56 μm/h
histogram
speed distribution μm/h

#### Supplementary Files

**Figure_S3-A: Localized analysis of oncostream formation in distinct zones. (a)** This figure presents a representative time-lapse scanning confocal image from **Movie #3** (main manuscript Figure 4), captured at the end of a 24-hour time-lapse, showing glioma cells and the division into three distinct zones (A, B, and C). This segmentation allows for a detailed zone-specific analysis of cell behavior during oncostream formation. **(b)** Angular velocity distribution, represented by angle θ, is analyzed for each zone to demonstrate the preferred movement direction of cells at different stages of oncostream development. The stages include early (0-6hr), mid (6-12hr), late (12-18hr), and final (18-24hr) periods, offering insights into how movement speed and directionality evolve over time in Zones A, B, and C. **(c)** The distribution of cell speeds and the average speed (measured in μm/h) for each time interval are calculated for Zones A, B, and C. This data provides a comprehensive view of the kinetic behavior of cells and their collective migration patterns during each stage of the time-lapse.

**Figure S3-B**

**Time-lapse 0-6hr**

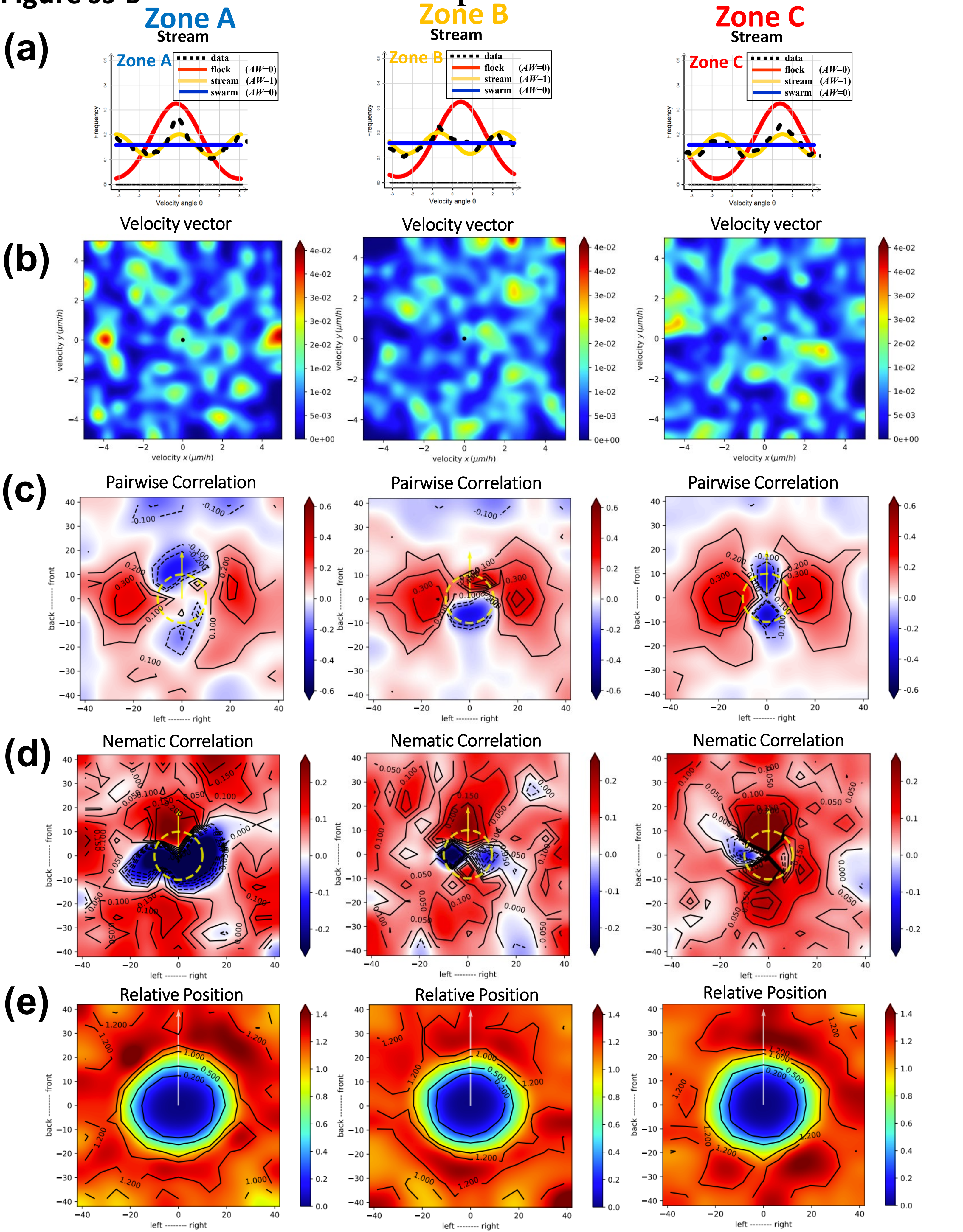

#### Supplementary Files

**Figure_S3-B: Extended localized analysis of oncostream dynamics in zones A, B, and C during the early stage (0-6hr). (a)** A likelihood analysis of frequency distributions in the early stage (0-6hr) reveals a coordinated, organized behavior in oncostream formation when characterized locally in distinct zones A, B, and C. **(b)** Heatmaps of velocity vector distributions for the early (0-6hr) phase of the time-lapse offer an in-depth view of the dynamic oncostream formation in zones A, B, and C. These visualizations facilitate the observation of alterations in movement speed and directionality across the different zones. **(c)** Pair-wise velocity correlations among neighboring cells are quantified for each zone, considering relative positions (xi - xj) and directional correlation (ωi to ωj). The axes represent cell positions and distances in micrometers, and a dotted yellow line indicates the average size of the central cell. This analysis illustrates the coordination of cell motility during early-stage oncostream development in each zone. **(d)** Heatmaps of nematic correlations, based on the proximity of adjacent cells, underscore the alignment and directionality of cell movements in Zones A, B, and C during the early stage (0-6hr). These heatmaps provide insight into the collective behavior and organization of cells in each specific zone. **(e)** The frequency of relative positioning between cell pairs in the early stage (0-6hr) is analyzed to estimate the probability of cell proximity, mapping spatial interactions and matrix-dependent dynamics. This analysis aids in understanding the spatiotemporal structural evolution of oncostreams within distinct zones A, B, and C, offering a detailed perspective on how cell organization unfolds over time.

**Figure S3-C**

### Time-lapse 6-12hr

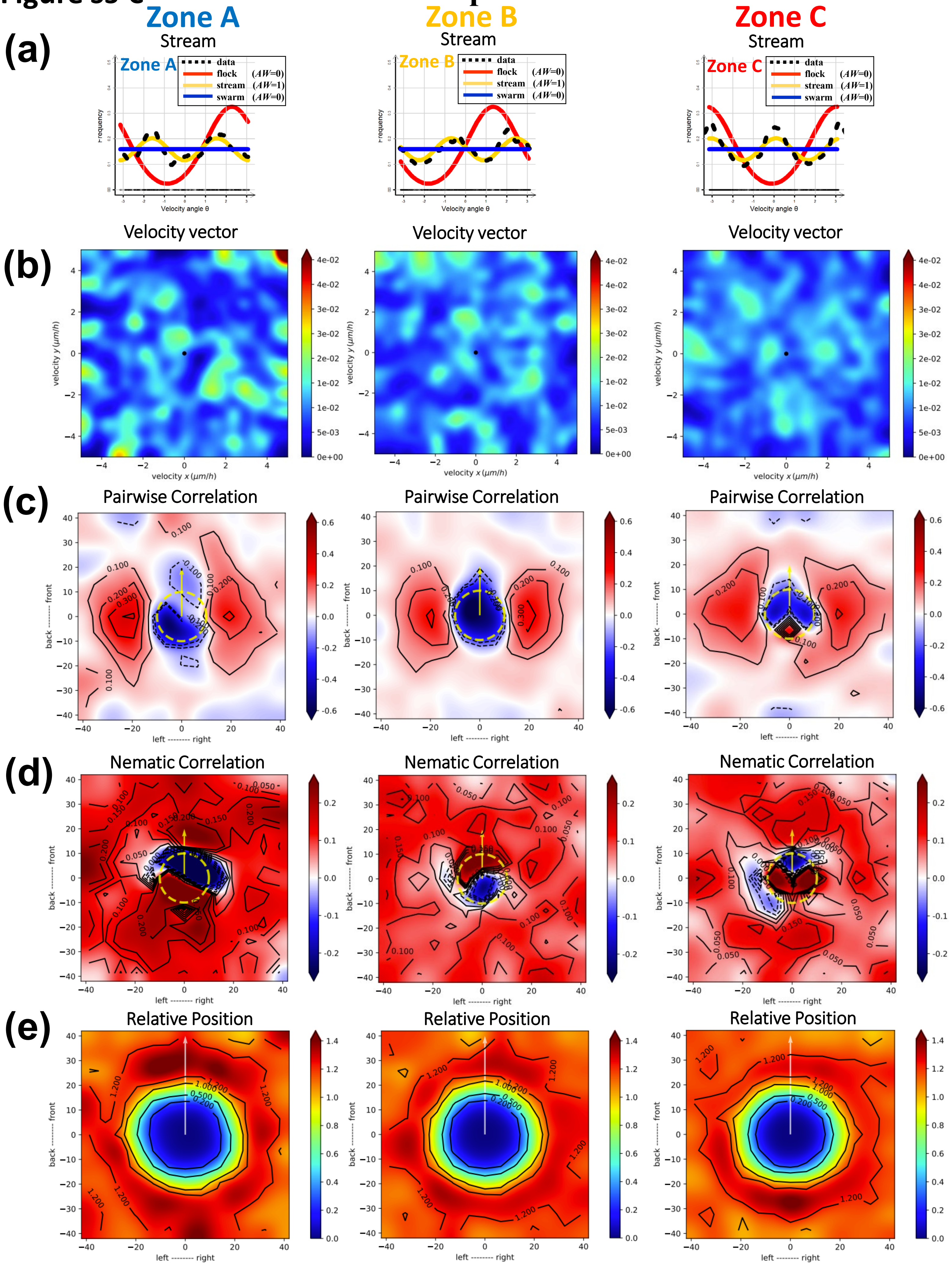

#### Supplementary Files

**Figure_S3-C: Extended localized analysis of oncostream dynamics in zones A, B, and C during the mid stage (6-12hr). (a)** Likelihood analysis of frequency distributions in the mid stage (6-12hr) shows a coordinated, organized behavior in oncostream formation across distinct zones A, B, and C. This highlights structured cell movement patterns during this critical phase of development. **(b)** Heatmaps of velocity vector distributions for the mid stage (6-12hr) provide a comprehensive view of dynamic oncostream formation in zones A, B, and C. These visualizations allow for the observation of changes in movement speed and directionality in each zone. **(c)** Pair-wise velocity correlations among neighboring cells are quantified within each zone, considering their relative positions ($x_i$ - $x_j$) and directional correlation ($\omega_i$ to $\omega_j$). The axes denote cell positions and distances in micrometers, with a dotted yellow line marking the average size of a central cell. This analysis illustrates the coordination of cell motility during the mid-stage of oncostream development. **(d)** Heatmaps of nematic correlations based on the proximity of adjacent cells highlight the alignment and directionality of cell movements in zones A, B, and C during the mid stage (6-12hr). These heatmaps provide insights into the collective behavior and organization of cells within each specific zone. **(e)** The frequency of relative positioning between cell pairs in the mid stage (6-12hr) is analyzed, estimating the likelihood of cell proximity to map spatial interactions and matrix-dependent dynamics. This analysis offers a detailed perspective on the spatiotemporal structural evolution of oncostreams within the distinct zones A, B, and C, elucidating how cell organization progresses over this time frame.

### Figure S3-D

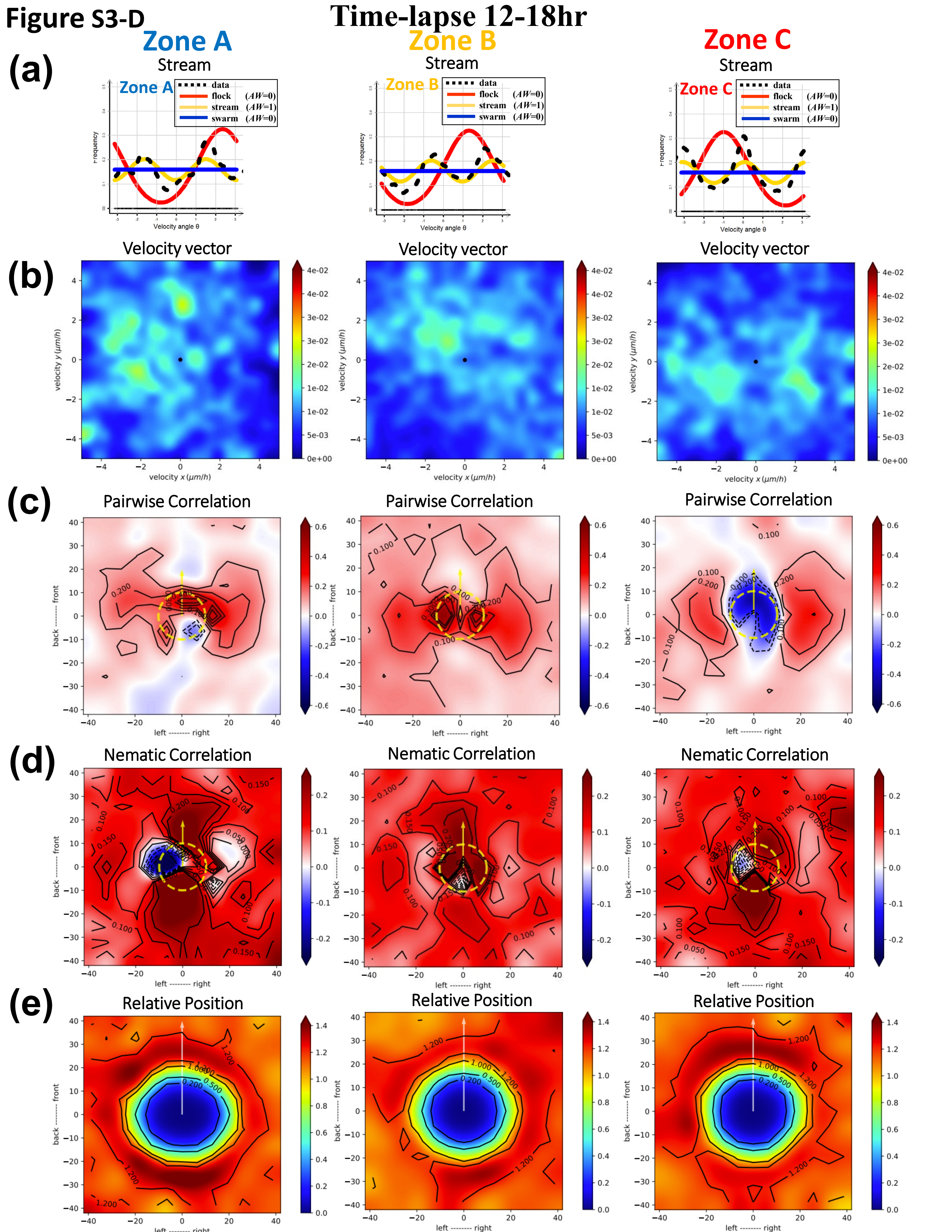

Time-lapse 12-18hr
Zone A
Zone B
Zone C
(a)
Stream
data
flock (AW=0)
stream (AW=1)
swarm (AW=0)
Frequency
Velocity angle θ
(b)
Velocity vector
velocity y (μm/h)
velocity x (μm/h)
(c)
Pairwise Correlation
back -------- front
left -------- right
(d)
Nematic Correlation
(e)
Relative Position

#### Supplementary Files

**Figure_S3-D: Extended localized analysis of oncostream dynamics in zones A, B, and C during the late stage (12-18hr). (a)** Likelihood analysis of frequency distributions in the late stage (12-18hr) reveals coordinated, organized behavior in oncostream formation across distinct zones A, B, and C. This emphasizes structured cell movement patterns in the advanced phase of development. **(b)** Heatmaps of velocity vector distributions for the late stage (12-18hr) offer an in-depth view of the dynamic oncostream formation in zones A, B, and C. These visualizations facilitate observations of changes in movement speed and directionality within each zone during this critical period. **(c)** Pair-wise velocity correlations among neighboring cells are quantified for each zone during the late stage, considering their relative positions (xi - xj) and directional correlation (ωi to ωj). The axes represent cell positions and distances in micrometers, with a dotted yellow line indicating the average size of a central cell. This analysis demonstrates the coordination of cell motility during this later stage of oncostream development. **(d)** Heatmaps of nematic correlations, based on the proximity of adjacent cells, underscore the alignment and directionality of cell movements in zones A, B, and C during the late stage (12-18hr). These heatmaps provide valuable insights into the collective behavior and organization of cells in each specific zone. **(e)** The frequency of relative positioning between cell pairs in the late stage (12-18hr) is evaluated, estimating the probability of proximity to map spatial interactions and matrix-dependent dynamics. This analysis provides a comprehensive view of the spatiotemporal structural evolution of oncostreams in the distinct zones A, B, and C, illustrating how cell organization continues to evolve in this later phase.

Figure S3-E

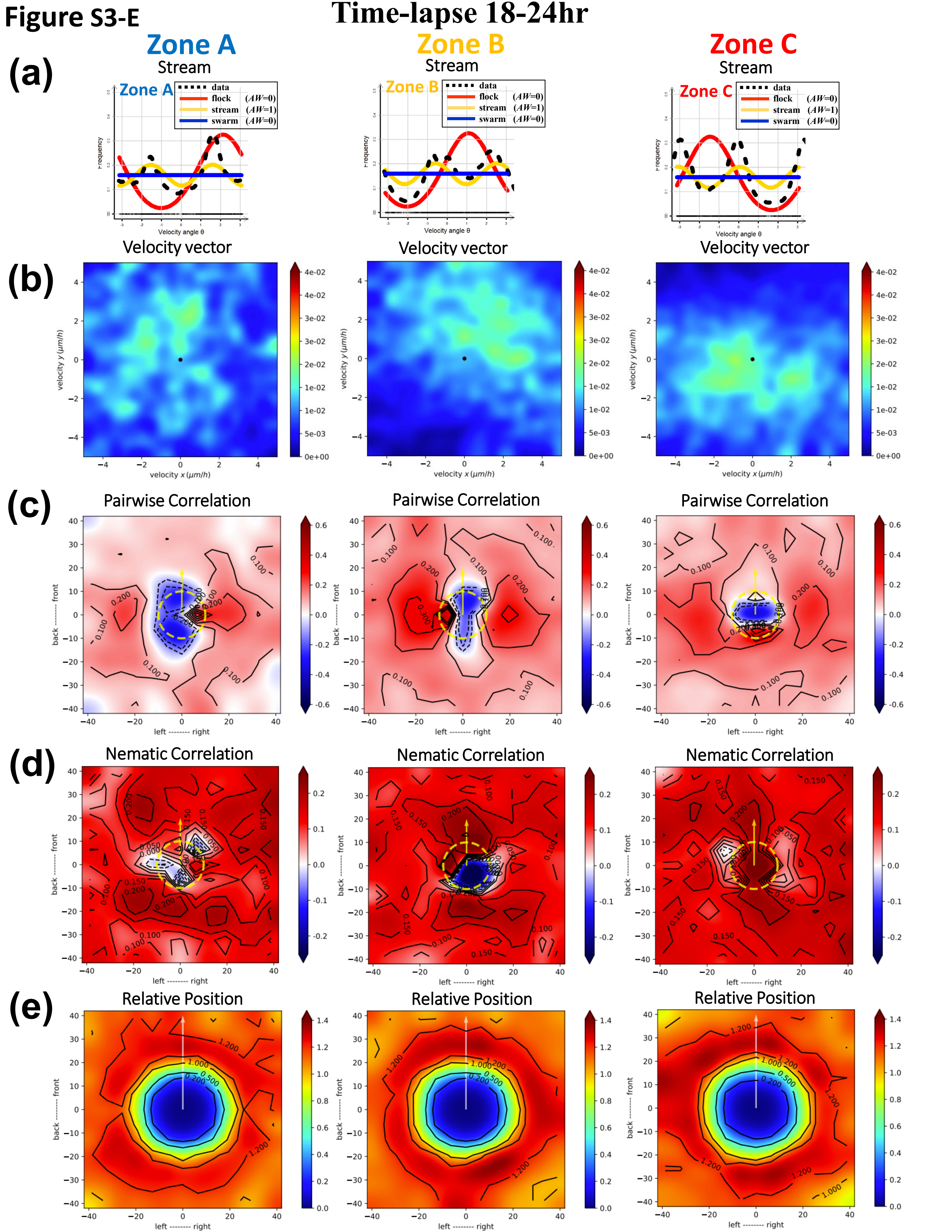
Time-lapse 18-24hr
Zone A
Zone B
Zone C
(a)
Stream
data
flock (AW=0)
stream (AW=1)
swarm (AW=0)
Frequency
Velocity angle θ
Velocity vector
(b)
velocity y (μm/h)
velocity x (μm/h)
(c)
Pairwise Correlation
(d)
Nematic Correlation
(e)
Relative Position
back -------- front
left -------- right

#### Supplementary Files

**Figure_S3-E: Extended localized analysis of oncostream dynamics in zones A, B, and C during the final stage (18-24hr). (a)** Likelihood analysis of frequency distributions in the final stage (18-24hr) demonstrates a sustained, coordinated behavior in oncostream formation within zones A, B, and C. This indicates the culmination of structured cell movement patterns as the oncostreams reach maturity. **(b)** Heatmaps of velocity vector distributions for this final stage offer a comprehensive view of the oncostream formation dynamics in zones A, B, and C. These visualizations highlight changes in movement speed and directionality, marking the evolution of cell behavior as the process concludes. **(c)** Pair-wise velocity correlations among neighboring cells are quantified in each zone during the final stage, considering their relative positions (xi - xj) and directional correlation (ωi to ωj). The axes denote cell positions and distances in micrometers, with a dotted yellow line illustrating the average size of a central cell. This analysis underscores the coordination and final adjustments in cell motility as the oncostreams fully develop. **(d)** Heatmaps of nematic correlations, based on the proximity of adjacent cells, reveal the alignment and directionality of cell movements in zones A, B, and C during the final stage. These heatmaps offer insights into the collective behavior and organization of cells, showcasing the final arrangement within each specific zone. **(e)** The frequency of relative positioning between cell pairs in the final stage (18-24hr) is analyzed, providing an estimation of cell proximity to map spatial interactions and matrix-dependent dynamics. This comprehensive analysis elucidates the final phase of the spatiotemporal structural evolution of oncostreams across the distinct zones A, B, and C, highlighting the culmination of cell organization patterns.

**Figure S4** 4-HAP *(Non-Muscle Myosin IIC)* do not dismantles oncostreams *in vitro*

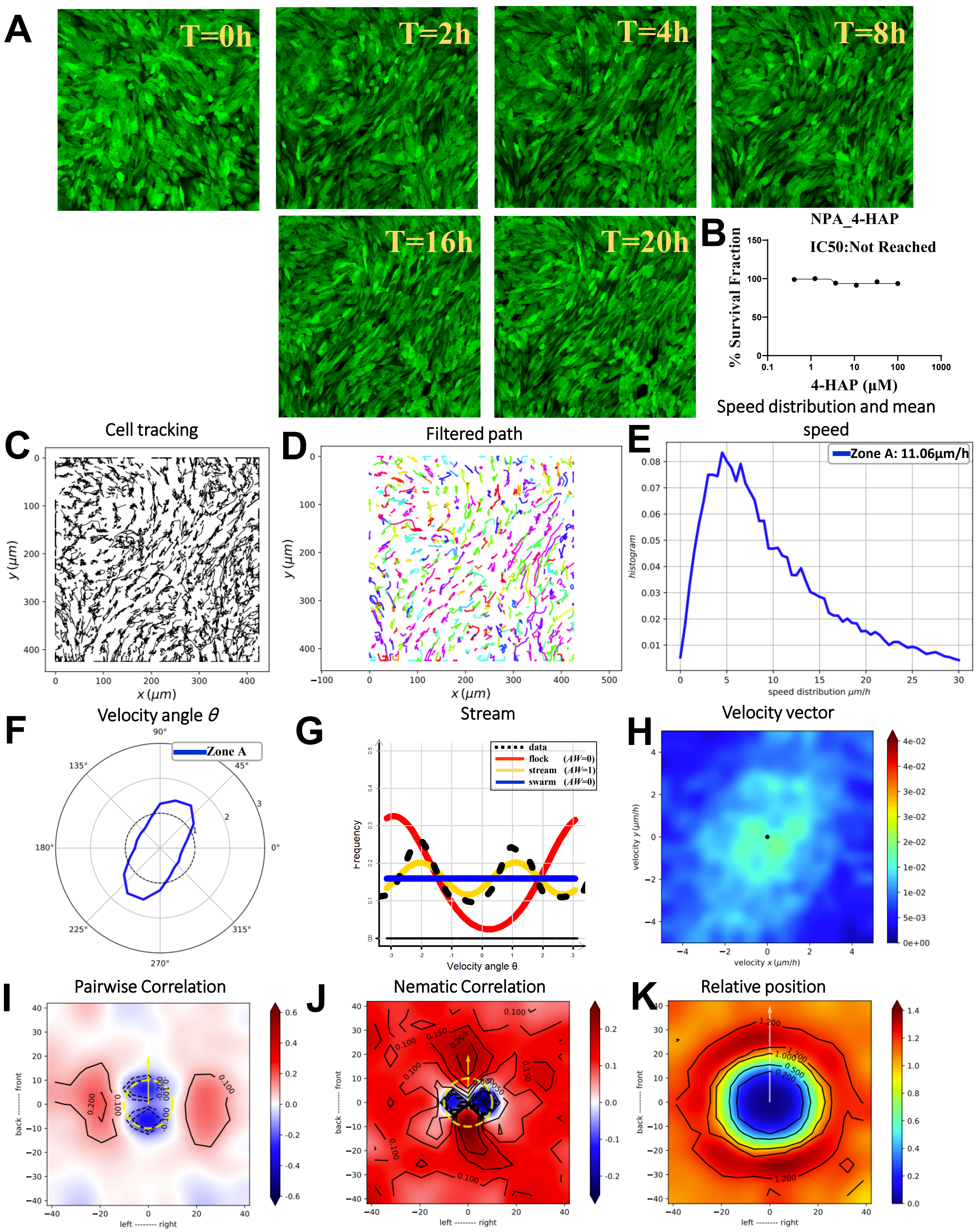

**Supplementary Files**

**Figure_S4: Effect of 4-HAP on oncostream structure.** This figure illustrates the impact of 4-HAP treatment (a non-muscle myosin IIC inhibitor) on oncostreams' structure, with dynamics monitored over a 20-hour period using confocal time-lapse imaging (refer to **Movie #13**). **(A)** Time-lapse images at 0, 2, 4, 8, 16, and 20 hours showcase cell organization within fully formed oncostreams treated with 4-HAP, highlighting the dynamic behavior of oncostreams. **(B)** Dose-response curves for NPA glioma cells treated with varying concentrations of 4-HAP (100 mM to 0.411 mM) are presented. IC50 values were not achieved. A concentration of 4 μM 4-HAP was selected based on the curve to ensure effective treatment without reaching toxic levels. **(C)** Cell trajectories were tracked using the TrackMate plugin in Fiji, illustrating movement patterns over time and the influence of 4-HAP treatment on these trajectories. **(D)** Cell trajectories were further filtered using a Gaussian kernel filter ($\sigma = 2$, stencil of 9 points) in a Julia-based script, highlighting movement patterns in the treated group. **(E)** The distribution of cell speeds and the mean speed were analyzed, showing no significant change in cell motility post-treatment (11.06 μm/h) compared to untreated conditions (11.57 μm/h). **(F)** Angular velocity distribution is depicted, illustrating directional movement. The angle $\theta$ indicates cells' preferred movement directions under 4-HAP treatment, showing intact oncostream structure during the treatment. **(G)** Non-parametric estimation and Akaike weight analysis of data distributions indicate that 4-HAP treatment maintains stream-like structures without significant variations in cell motility. **(H)** Heatmaps demonstrate velocity vector distributions throughout the time-lapse period, similar oncostream dynamics in cells treated with 4-HAP compared to the untreated group. **(I)** Pair-wise velocity correlations among neighboring cells were assessed. The analysis considered their relative positions ($x_i - x_j$) and directional correlation ($\omega_i$ to $\omega_j$), with axes representing cell positions and distances in μm. A dotted yellow line indicates the average size of the centered cell. **(J)** Nematic correlation heatmaps show movement alignment, with red indicating positive (synchronous) correlation and blue indicating negative (opposite direction) correlation. **(K)** The spatial distribution and interaction dynamics (relative position) influenced by the treatments are quantified, mapping the frequency and probability distributions of cells in proximity.

**Figure S5**

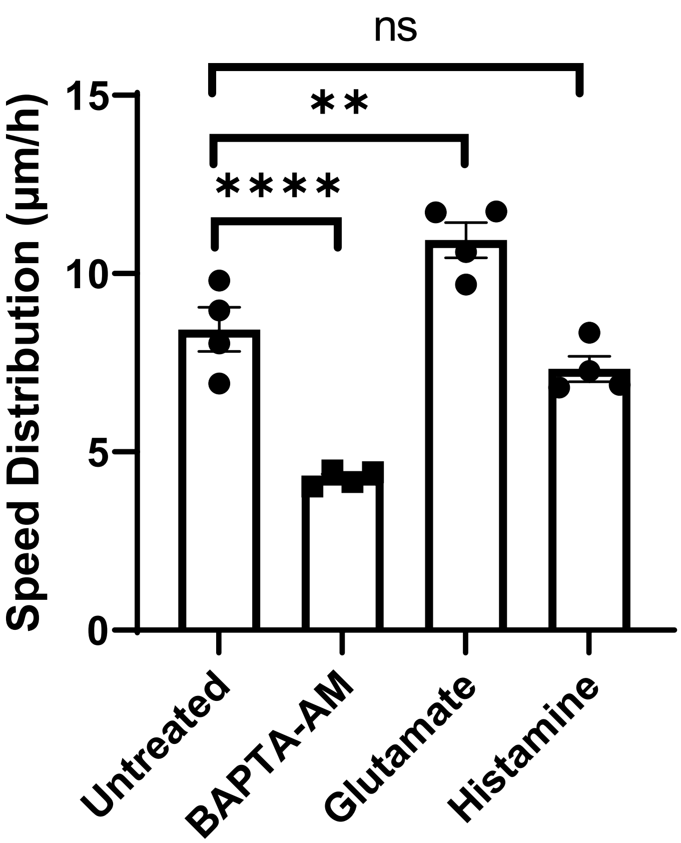
ns
**
****
15
10
5
0
Speed Distribution (µm/h)
Untreated
BAPTA-AM
Glutamate
Histamine

**Supplementary Files**

**Figure_S5: Speed Distribution in Response to Neurotransmitter Agonist and Antagonist Treatment.** This figure examines the impact of various neurotransmitter agonist and antagonist treatments on the collective migration kinetics of cells. The distribution of cell speeds and mean speed (measured in μm/h) were calculated to gain insights into the kinetic behavior of cells under different treatment conditions. Four time-lapse series were analyzed for each treated and control group to ensure the strength of the data. Statistical significance was determined using a non-parametric t-test, with each dot on the bar graph representing the speed distribution from one time-lapse series. The data are presented as mean with SEM. Significance levels are denoted as follows: ****$p < 0.0001$, ***$p < 0.001$, **$p < 0.01$, *$p < 0.05$, with $p > 0.05$ considered non-significant.

Figure S6

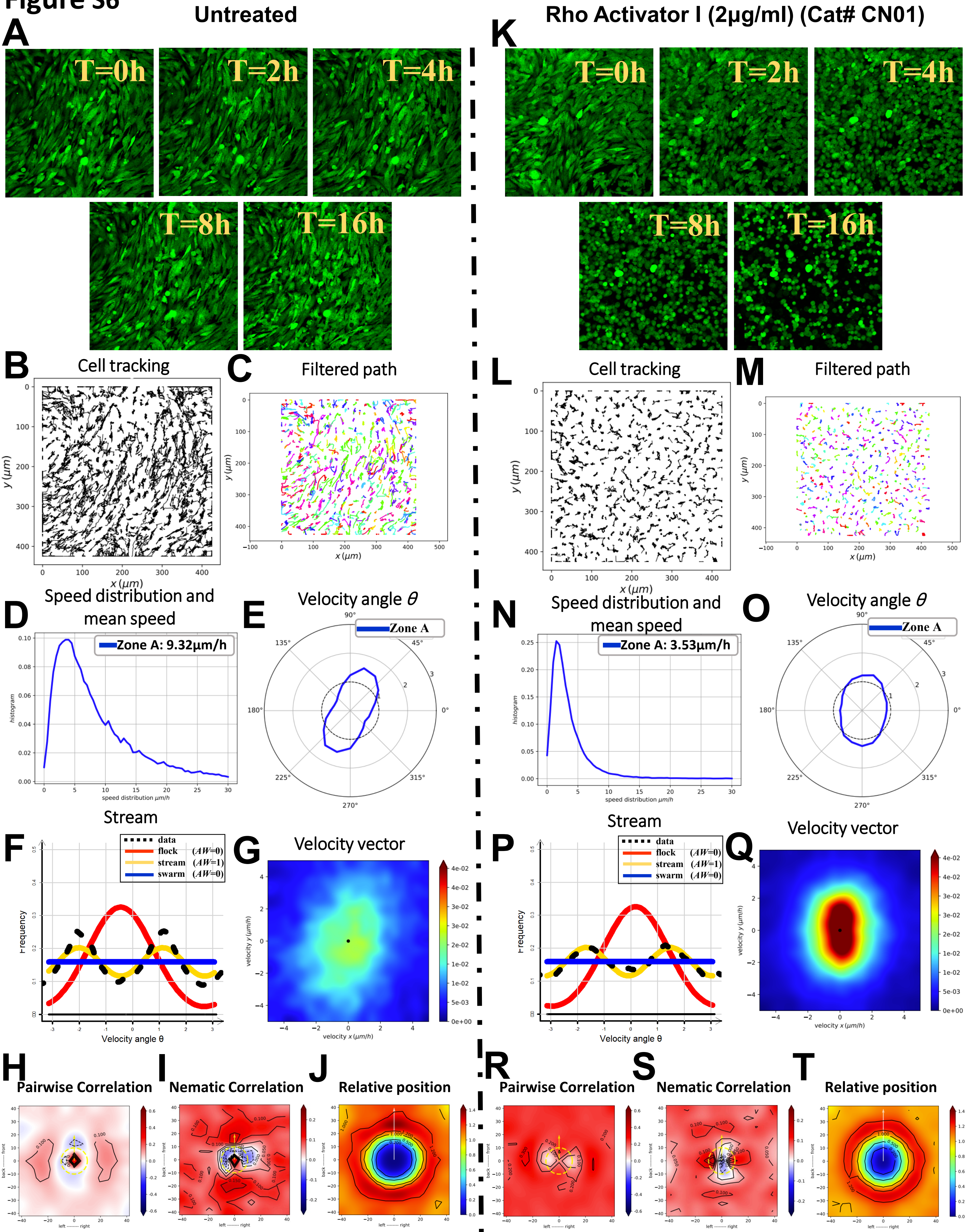

Untreated
Rho Activator I (2μg/ml) (Cat# CN01)
A
K
T=0h
T=2h
T=4h
T=8h
T=16h
B
Cell tracking
C
Filtered path
L
M
D
Speed distribution and mean speed
Zone A: 9.32μm/h
E
Velocity angle θ
Zone A
N
Zone A: 3.53μm/h
O
F
Stream
data
flock (AW=0)
stream (AW=1)
swarm (AW=0)
Frequency
Velocity angle θ
G
Velocity vector
P
Q
H
Pairwise Correlation
I
Nematic Correlation
J
Relative position
R
S
T
y (μm)
x (μm)
histogram
speed distribution μm/h
velocity y (μm/h)
velocity x (μm/h)

#### Supplementary Files

**Figure S6: Impact of Rho-Activator I on Oncostream Formation and Dynamics.** This figure illustrates the influence of Rho-Activator I on the formation and dynamics of oncostreams. Cellular behavior was recorded over a 16-hour period using confocal time-lapse imaging (refer to **Movie #14** for untreated and **Movie #15** for Rho-Activator I treatment). **(A, K)** Comparative time-lapse images at 0, 2, 4, 8, and 16 hours display cell organization in untreated conditions versus those treated with Rho-Activator I, highlighting the impact on oncostream integrity. **(B, L)** The TrackMate plugin in Fiji software was utilized to track individual cell paths, shedding light on the movement patterns and the effects of Rho-Activator I treatment on cell trajectories. **(C, M)** Cell trajectories for both untreated and treated groups were smoothed using a Gaussian kernel filter ($\sigma = 2$, stencil of 9 points) implemented in a Julia-based script, allowing for a clearer visualization of movement patterns. **(D, N)** Analysis of cell speed distributions revealed a significant decrease in motility following Rho-Activator I treatment, with mean speeds dropping from 9.32 μm/h in untreated cells to 3.53 μm/h in treated cells. **(E, O)** Angular velocity distribution plots were generated to analyze the directionality of cell movement, with the angle $\theta$ indicating the preferred direction of movement under both untreated and treated conditions. **(F, P)** Non-parametric estimation of data distributions and Akaike weight (AW) analysis suggested that Rho-Activator I-treated cells exhibit a streaming behavior. However, visually, they appear disorganized compared to untreated cells, which retain stream-like structures both visually and in likelihood analysis. **(G, Q)** Velocity vector distribution heatmaps for the entire time-lapse period illustrate the contrasting dynamics between untreated and treated cells, with respect to oncostream behavior. **(H, R)** Pair-wise velocity correlations among neighboring cells were calculated by considering their relative positions ($x_i - x_j$) and directional correlation ($\omega_i$ to $\omega_j$), with a dotted yellow line representing the average size of a centered cell. **(I, S)** Heatmaps of nematic correlations in proximity to neighboring cells provide insights into the alignment of movement, with red indicating synchronous movement and blue indicating opposite directions. **(J, T)** Spatial distribution and interaction dynamics influenced by Rho-Activator I treatment were quantified, estimating the frequency and probability of cell proximities to assess changes in cell-cell interactions and relative position.

### Figure S7

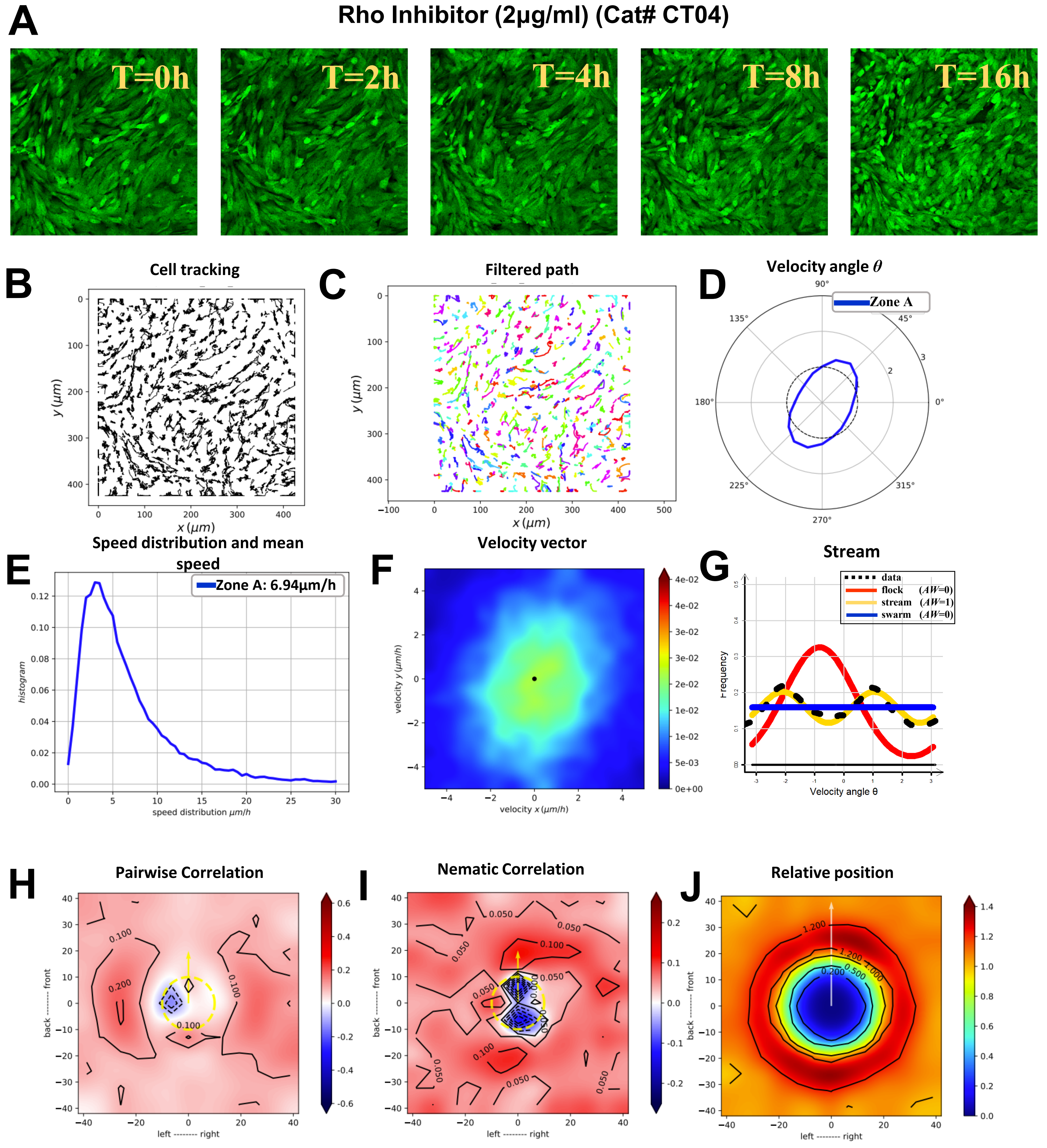

A
Rho Inhibitor (2µg/ml) (Cat# CT04)
T=0h
T=2h
T=4h
T=8h
T=16h
B
Cell tracking
y (µm)
x (µm)
C
Filtered path
y (µm)
x (µm)
D
Velocity angle θ
Zone A
E
Speed distribution and mean speed
Zone A: 6.94µm/h
histogram
speed distribution µm/h
F
Velocity vector
velocity y (µm/h)
velocity x (µm/h)
G
Stream
data
flock (AW=0)
stream (AW=1)
swarm (AW=0)
Frequency
Velocity angle θ
H
Pairwise Correlation
back -------- front
left -------- right
I
Nematic Correlation
back -------- front
left -------- right
J
Relative position
back -------- front
left -------- right

#### Supplementary Files

**Figure S7: Impact of Rho-Inhibitor on Oncostream Formation and Dynamics.** This figure illustrates the effects of Rho-Inhibitor on the formation and dynamics of oncostreams. Cellular behavior was observed over a 16-hour period using confocal time-lapse imaging (refer to **Movie #16** for Rho-Inhibitor treatment). **(A)** Time-lapse images taken at 0, 2, 4, 8, and 16 hours illustrate the organization of cells treated with Rho-Inhibitor, demonstrating changes in oncostream integrity. **(B)** The TrackMate plugin in Fiji software was employed to track individual cell paths, providing insight into movement patterns and the impact of Rho-Inhibitor treatment on these trajectories. **(C)** Cell trajectories of the Rho-Inhibitor treated group were refined using a Gaussian kernel filter ($\sigma = 2$, stencil of 9 points) with a Julia-based script, facilitating an enhanced visualization of movement patterns. **(D)** Angular velocity distribution plots were generated to evaluate the directionality of cell movement, with the angle $\theta$ indicating the preferred direction of movement in the Rho-Inhibitor treated condition. **(E)** Cell speed distribution analysis showed a significant reduction in motility following Rho-Inhibitor treatment, with average speeds decreasing from 9.32 μm/h in control cells to 6.94 μm/h in treated cells. **(F)** Heatmaps of velocity vector distributions throughout the time-lapse period highlight the altered dynamics of oncostreams under Rho-Inhibitor treatment. **(G)** Non-parametric estimation and Akaike weight (AW) analysis of data distributions indicated that Rho-Inhibitor-treated cells maintain streaming behavior. **(H)** Pair-wise velocity correlations among neighboring cells was assessed, considering their relative positions ($x_i$ - $x_j$) and calculating the directional correlation ($\omega_i$ to $\omega_j$). A dotted yellow line marks the average size of a centered cell for reference. **(I)** Heatmaps of nematic correlations based on the proximity of neighboring cells assess the alignment of movement, with red indicating synchronous movement and blue indicating movement in opposite directions. **(J)** The spatial distribution and interaction dynamics influenced by Rho-Inhibitor treatment were quantified. The frequency and probability of cell proximities were estimated to evaluate changes in cell-cell interactions and relative position.

#### DESCRIPTION OF MOVIES

**Manuscript: Modeling Glioma Oncostreams In Vitro: Spatiotemporal Dynamics of their Formation, Stability, and Disassembly.**

Movies are available at Zenodo (https://doi.org/10.5281/zenodo.10381916).

**Movie 1: Low Density Oncostream Dynamics:** Demonstrates the formation and dynamics of oncostreams in GFP+ NPA glioma cells at low seeding density (1 × 10^5 cells), over 24 hours using time-lapse confocal imaging.

**Movie 2: High Density Oncostream Dynamics:** Explores the effect of high cell seeding density (2 × 10^5 cells) on the formation and behavior of oncostreams in GFP+ NPA glioma cells, analyzed over 24 hours.

**Movie 3: Spatiotemporal Progression of Oncostreams:** Showcases the sequential self-formation and developmental stages of oncostreams over a 24-hour period.

**Movie 4: Collagenase Impact on Oncostreams:** Description: Illustrates the disassembly of aggressive and malignant oncostreams with 15 U/ml collagenase treatment over 15 hours.

**Movie 5: TC-I-15 Inhibition of Oncostream Formation:** Details the impact of TC-I-15, an integrin antagonist, on adhesion and oncostream formation in glioma cells, recorded over 20 hours.

**Movie 6: Cytochalasin D Disruption of Oncostreams:** Captures the effect of Cytochalasin D on oncostream formation by inhibiting actin polymerization, observed over a 1-hour period with rapid imaging intervals.

**Movie 7: Myosin II Inhibition in Oncostreams with p-nitro Blebbistatin:** Presents the influence of p-nitro Blebbistatin on oncostream formation by inhibiting myosin II, documented over 1 hour with dynamic cellular responses.

**Movie 8: Control - Oncostream Formation without BAPTA-AM:** Demonstrates oncostream formation and dynamics without BAPTA-AM treatment, monitored over 16 hours.

**Movie 9: BAPTA-AM (Calcium Modulation) Impact on Oncostream Formation:** Reveals the effects of BAPTA-AM treatment on oncostream dynamics, observed over a 16-hour period.

**Movie 10: Glutamate Influence on Oncostreams:** Shows the effects of glutamate treatment on oncostream formation and dynamics, monitored over 16 hours.

**Movie 11: Histamine Impact on Oncostream Dynamics:** Demonstrates the influence of histamine on the formation and behavior of oncostreams, captured over a 16-hour period.

**Movie 12: Oncostream Formation on Non-Laminin Coated Surfaces:** Illustrates the compromised organization of oncostreams on poly-D-lysine coated dishes without laminin, observed over 45 hours.

**Movie 13: 4-HAP Treatment Effect on Oncostreams:** Shows the impact of 4-HAP treatment on the structure of oncostreams, monitored over 20 hours.

**Movie 14: Rho-Activator I Untreated Control:** Presents the formation and dynamics of oncostreams without Rho-Activator I treatment, recorded over 16 hours.

**Movie 15: Rho-Activator I Influence on Oncostreams Dynamics:** Details the effects of Rho-Activator I on oncostream formation and dynamics, observed over a 16-hour period.

**Movie 16: Rho-Inhibitor Effect on Oncostreams Dynamics:** Demonstrates the impact of Rho-Inhibitor on the formation and behavior of oncostreams, monitored over 16 hours.
